# Parkinson’s disease-associated alterations in DNA methylation and hydroxymethylation in human brain

**DOI:** 10.1101/2024.05.21.595193

**Authors:** Juliana I. Choza, Mahek Virani, Nathan C. Kuhn, Marie Adams, Joseph Kochmanski, Alison I. Bernstein

**Author notes:** **Corresponding Author:** Alison Bernstein, Environmental and Occupational Health Sciences Institute, Ernest Mario School of Pharmacy, Rutgers University 170 Freylinghuysen Rd, Piscataway, NJ 08854. These authors contributed equally to this work (Juliana I. Choza, Mahek Virani).

## Abstract

Epigenetic mechanisms are mediators of interactions between aging, genetics, and environmental factors in sporadic Parkinson’s disease (PD). Multiple studies have explored the DNA modifications in PD, but few focus on 5-hydroxymethylcytosine (5-hmC), which is important in the central nervous system and sensitive to environmental exposures. To date, studies have not differentiated between 5-methylcytosine (5-mC) and 5-hmC or have analyzed them separately. In this study, we modeled paired 5-mC and 5-hmC data simultaneously. We identified 108 cytosines with significant PD-associated shifts between these marks in an enriched neuronal population from PD postmortem parietal cortex, within 83 genes and 34 enhancers associated with 67 genes. These data potentially link epigenetic regulation of genes related to LRRK2 and endolysosomal sort (*RAB32* and *AGAP1*), and genes involved in neuroinflammation, the inflammasome, and neurodevelopment with early changes in PD and suggest that there are significant shifts between 5mC and 5hmC associated with PD in genes not captured by standard methods.

## Introduction

An estimated 5-10% of Parkinson’s disease (PD) cases are familial and caused by monogenically inherited mutations, while the remaining ∼90% of sporadic cases (sPD) are likely due to a complex interaction between age, genes, and environmental factors ^1–3^. While the relative contribution of genetic and environmental risk factors in the etiology of sPD is debated, it is well documented that they play critical roles in the large majority of PD cases. Epigenetic mechanisms have emerged as critical mediators of the complex interactions between aging, genetics, and the environment because they are dynamic with age, sensitive to the environment, and regulate gene expression throughout the lifespan ^4–6^. Evidence for the role of epigenetic regulation in PD has been building, particularly for DNA modifications ^7–10^.

5-methylcytosine (5mC), the addition of a methyl group to the 5′-carbon of cytosine, is one of the most well-studied epigenetic marks. 5-hydroxymethylcytosine (5hmC) is formed via oxidation of 5mC by ten-eleven translocation (TET) enzymes and is a stable, independent epigenetic mark that has its highest levels in the brain, recruits a distinct set of DNA binding proteins from 5mC, differs in its genomic distribution in the brain compared 5mC, and is enriched in transcriptionally active gene bodies in the nervous system, suggesting a specific regulatory role for 5hmC in the brain ^11^. Thus, 5hmC is now thought to be particularly important in gene regulation in the brain, particularly in the response to environmental exposures and neurotoxicants ^12, 13^. However, most studies of DNA modifications in PD brain have relied on bisulfite (BS) conversion, which cannot distinguish between 5mC and 5hmC ^14, 15^.

Recently, studies have begun to explore links between 5hmC and PD ^16–18^. First, rare variants in *TET1* were associated with an increased risk of PD in a Chinese PD cohort ^16^. Second, a targeted analysis of DNA modifications within known enhancers in human postmortem prefrontal cortex identified epigenetic disruption of an enhancer targeting the *TET2* gene in PD patients ^18^. This study also performed hydroxymethylated DNA immunoprecipitation-sequencing (hMeDIP-Seq) in prefrontal cortex and found that PD-associated-hydroxymethylated peaks were enriched in gene bodies, promoters, and enhancers. Third, a small study in human postmortem substantia nigra (SN) used hMe-Seal, a selective chemical labeling method, and identified thousands of differentially hydroxymethylated regions in genes related to CNS and neuronal differentiation, neurogenesis, and development and maintenance of neurites and axons, although the widespread neurodegeneration in the SN by the time of PD diagnosis complicates interpretation of these results ^17^. Regardless, taken together, these initial studies support a role for 5hmC in regulation of expression of genes important for PD pathogenesis and indicate that additional research is warranted.

In our previous study, we performed a neuron-specific epigenome-wide association study (EWAS) with the Illumina EPIC BeadChip array paired with BS conversion using enriched neuronal nuclei from human postmortem parietal cortex obtained from the Banner Sun Health Research Institute Brain Bank ^19^. We identified largely sex-specific PD-associated changes in DNA modification in 434 unique genes, including genes previously implicated in PD, including *PARK7* (DJ-1), *SLC17A6* (VGLUT2), *PTPRN2* (IA-2β), and *NR4A2* (NURR-1), as well as genes involved in developmental pathways, neurotransmitter packaging and release, and axon/neuron projection guidance. However, we could not differentiate between 5mC and 5hmC because we used BS conversion.

Here, we report the results of an EWAS of 5hmC and 5mC in enriched neurons from PD brain using our recently proposed method for reconciling base-pair resolution 5mC and 5hmC data ^20^. This method considers 5mC and 5hmC as paired data since these marks are biologically and statistically dependent on each other, and has been used for EWAS in human cohorts exposed to lead and PFAS ^21, 22^. We utilized additional DNA isolated from the same samples and performed oxidative BS (oxBS) conversion paired with the Illumina EPIC BeadChip array to specifically measure 5mC ^23, 24^. oxBS adds an oxidation step with potassium perruthenate (KRuO_4_) that specifically oxidizes 5hmC, forming 5-formylcytosine, prior to BS conversion. BS then deaminates 5mC only, but not 5-formylcytosine, such that C and 5hmC are read as thymine, while only the 5mC is read as cytosine, providing a readout of “true” methylation. Subsequent comparison of BS and oxBS results allows estimation of 5hmC. To our knowledge, this is the largest epigenome-wide analysis of 5hmC to date in neurons enriched from PD post-mortem brain at single base pair resolution.

## Methods

### Human brain tissue

De-identified tissue samples from control (n=50) and sPD (n=50) human brain samples were obtained from archival human autopsy specimens provided by the Banner Sun Health Research Institute (BSHRI), using BSHRI’s approved institutional review board (IRB) protocols. Further details about the BSHRI’s brain samples and sample selection are available in a previous publication ^25^. We selected PD patients with mid-stage disease (Braak stage = II–III), as defined by Lewy pathology. The cohort of control brains consisted of patients who died from non-neurologic causes and whose brains had no significant neurodegenerative disease pathology. For each subject (N=100), parietal cortex was obtained. This region develops pathology late in PD; in mid-stage PD and is expected to still have robust populations of neurons (unlike the substantia nigra, where neuron loss occurs early in disease), providing an avenue to investigate pre-pathological changes in gene regulation.

### Magnetic-activated cell sorting

NeuN-positive (NeuN^+^) nuclei were enriched from 100 mg of flash frozen parietal cortex tissue using a two-stage magnetic-assisted cell sorting (MACS) method as previously described ^19^.

First, 100 mg of frozen tissue was briefly thawed on ice and homogenized in a 2 mL, 1.4 mm ceramic bead tube (Thermo Fisher Scientific, Cat. # 15-340-153) with 1 mL of Nuclear Extraction Buffer (NEB) for 10 sec at 4 m/s. NEB consisted of 0.32M sucrose, 0.01M Tris-HCl pH 8.0, 0.005M CaCl_2_, 0.003M MgCl_2_, 0.0001M EDTA, and 0.1% Triton X-100, up to a stock volume of 1 L using water. Immediately prior to use, 0.001M DTT was added to NEB. Homogenized samples were loaded into a 13 mL ultracentrifuge tube (BeckmanCoulter, Cat. # 331372) with 4 mL of NEB. Using a glass pipette, 7 mL of sucrose solution was pipetted down the side of each sample tube to create a sucrose gradient. Sucrose solution consisted of 1.8M sucrose, 0.01M Tris-HCl pH 8.0, 0.003M MgCl_2_, up to a stock volume of 1 L using water.

Immediately prior to use, 0.001M DTT was added to NEB. After addition of sucrose, samples were spun at 4°C, 24,000 rpm in the Sorvall Wx+ Ultracentrifuge in a swing bucket rotor (TH-641). Once the centrifugation was complete, the supernatant and debris layer found at the concentration gradient were both removed with the use of a vacuum, while being careful not to disturb the pellet containing the nuclei at the bottom of the tube. Next, 1 mL of primary antibody (anti Neun 488 – Millipore, Cat. # MAB377X) in MACS buffer was added to each nuclei pellet and placed on ice for 10 minutes. MACS buffer consisted of 0.5% Bovine Serum Albumin solution (Sigma-Aldrich, Cat. # A1595) in PBS pH 7.2 (Gibco, Cat. # 20012-027). Samples were then mechanically pipetted up and down 10-15 times to completely dissolve the nuclei pellet within the primary antibody-MACS buffer solution. This solution of nuclei was then transferred to a 2 mL tube and incubated for 60 minutes at 4°C. After incubation, 40 µL of MACS Microbeads (anti-mouse IgG Microbeads - Miltenyi, Cat. # 130-048-401) were added to each sample. Samples were then inverted 4-5 times and incubated at 4°C for 30 minutes. After incubation, nuclei were centrifuged at 300 x g for 10 minutes. Supernatant was then removed, and the nuclei were resuspended in 2 mL of MACS buffer and transferred to a MACS MS column (MS Separation columns – Miltenyi, Cat. # 130-042-201) that was pre-washed with MACS buffer and attached to the Miltyeni OctoMACS™ Separator. Positive selection of NeuN^+^ cells was then performed according to the standard MACS MS Columns protocol available from Miltenyi Biotec. After the first round of magnetic separation, NeuN^+^ nuclei were run through a separate, second MACS MS column to maximize cell type enrichment. To validate our methods, a subset of isolated nuclei was analyzed for flow cytometry on a CytoFlex S (Beckman Coulter), and data were analyzed using FlowJo V10, as reported in our previous study^19^. Percent positivity for each sample was defined as the percentage of events in the NeuN-A488+ gate, divided by the total number of events identified as Nuclei. The average proportion of neurons was estimated to be 83.8% across all samples.

### DNA extraction

DNA was isolated from enriched NeuN^+^ nuclei using the Qiagen QIAamp DNA MicroKit (Cat. # 56304) as previously described with some modifications to maximize yield ^19^. Given that samples were already dissociated during nuclei isolation, the sample lysis and incubation steps of the QIAamp DNA Micro Kit protocol were removed. Instead, 20 µL of proteinase K were added directly to each MACS eluate. Samples were then vortexed for 15 seconds and incubated at room temperature for 15 minutes. In addition, the optional carrier RNA was added to Buffer AL, the incubation time after addition of 100% ethanol was increased to 10 minutes, the incubation time for the elution buffer was increased to 5 minutes, and the final elution step was repeated using 10 mM Tris-HCl pH 8.0.

### Oxidative bisulfite treatment and EPIC arrays

Intact genomic DNA yield was quantified by Qubit fluorometry (Life Technologies). Cleanup and preparation of DNA, and all steps for the EPIC bead chip protocol were performed as previously described per the manufacturer’s protocol, with additional steps for oxBS reactions ^19^.

Bisulfite conversion was performed on 500 ng genomic DNA using the TrueMethyl Array kit (Cambridge Epigenetix). oxBS conversion was performed on 1 µg genomic DNA using the TrueMethyl Array kit (Cambridge Epigenetix). While the recommended amount of 500 ng DNA was sufficient for BS conversions, this was insufficient for oxBS, likely due to the additional oxidation step. These reactions were initially carried out on the same batch of DNA with 500 ng input for both reactions. However, with this input amount, all oxBS reactions failed to hybridize to the array. Thus, the oxBS reactions were run on a new batch of DNA isolations.

All conversion reactions were cleaned using SPRI-bead purification and eluted in Tris buffer. Following elution, BS-and oxBS-converted DNA was denatured and processed through the EPIC array protocol. The EPIC array contains ∼850,000 probes that query DNA methylation at CpG sites across a variety of genomic features, including CpG islands, RefSeq genic regions, ENCODE open chromatin, ENCODE transcription factor binding sites, and FANTOM5 enhancer regions. To perform the assay, converted DNA was denatured with 0.4 N sodium hydroxide. Denatured DNA was then amplified, hybridized to the EPIC bead chip, and an extension reaction was performed using fluorophore-labeled nucleotides per the manufacturers protocol. Array BeadChips were scanned on the Illumina iScan platform.

### EPIC array data processing of oxBS data

IDAT files were imported into R and processed using an in-house bioinformatics pipeline that utilizes the following packages: *minfi* (version 1.48.0) for importing data, quality control, and dye bias correction, *ChAMP* (version 2.32.0) to perform singular value decomposition (SVD) to identify covariates for modeling, *posibatch* (version 1.0) for batch correction, and *ENmix* (version 1.38.01) to verify performance of control probes and perform a maximum likelihood estimate (MLE) of paired bisulfite and oxidative bisulfite, as previously described (Supplementary File 1) ^19, 20, 24, 26–30^. After QC, eleven female samples and six male samples were removed due to a high level (>10%) of failed probes, leaving 57 male (28 PD, 29 control) and 26 female (13 PD, 13 control) samples. One additional male sample was excluded because it was excluded in our previous BS-only analysis, leaving 56 male samples (27 PD, 29 control) (47).

Failed probes (53,305) were removed from remaining samples when detection p-value was > 0.01 in more than 5% of samples. Cross-reactive probes and probes containing SNPs (93,527) were masked based on previous identification ^31^.

We continued with male samples only due to the small sample size of the remaining female samples. Data for the included samples are summarized in Table 1 and full metadata is in Supplementary File 2. Although this study has a sample size smaller than recent recommendations published by Mansell et al for BS-only-based studies, our subject selection experimentally controls for multiple variables, including sex, race, and age, which were included as covariates in their estimates ^32^. This increases our statistical power relative to the proposed minimum sample size, but future studies should include larger cohorts or be combined with publicly available data.

**Table 1:** Cohort characteristics of included samples. Data includes disease status, age at death in years, postmortem interval (PMI) in hours, and race of samples remaining after QC. Seven male samples and eleven female samples were removed during quality control and pre-processing, leaving 56 male and 26 female samples.

|  |  | Male (n = 56) |  |
| --- | --- | --- | --- |
| Variables | | Mean ± SD or | Range |
| *Disease Status* |  |  |  |
|  | Control | 29 (51.8%) |  |
|  | Parkinson's disease | 27 (48.2%) |  |
| *Age at Death* |  |  |  |
| | Control | 79.3 ± 9.1 | 53-93 |
| | Parkinson's disease | 79.4 ± 7.1 | 64-91 |
| *PMI* |  |  |  |
| | Control | 3.25 ± 0.81 | 2.16-5.5 |
| | Parkinson's disease | 3.27 ± 0.83 | 1.83-4.92 |
| *Race* |  |  |  |
|  | White | 55 (98.2%) |  |
|  | Unidentified | 1 (1.8%) |  |

5hmC β values (β_hmC_) were estimated by pairing oxBS β values with BS β values using the maximum likelihood estimate function (oxBS.MLE) from the *ENmix* package, which returns true methylation β values (β_mC_) and estimates β_hmC_. Density plots of raw BS and oxBS β values, as well as MLE-corrected β_mC_ and β_hmC_ values, are shown in Figure 1A,B.

**Figure 1:**
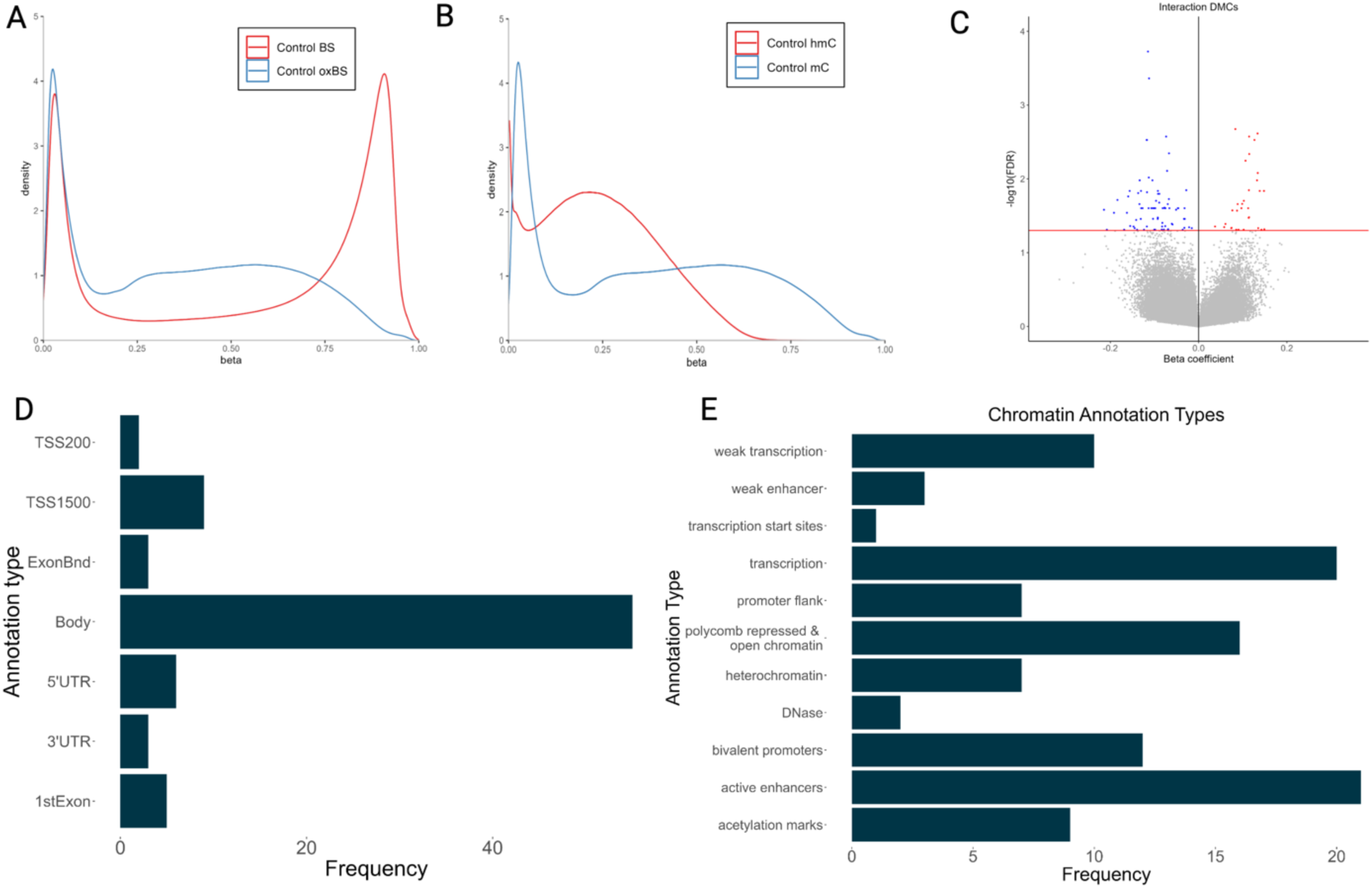
Identification of PD-associated iDMCs. Desnity plots of (A) raw BS and oxBS β-values and (B) MLE-corrected β_mC_ and β_hmC_ (B). (C) Volcano plot of iDMCs derived from MLE-corrected β values input into differential modification analysis (FDR < 0.05). Red indicates an increased interaction term; blue indicates a decreased interaction term. (D,E) Frequency histograms showing annotations of significant iDMCs for (D) intragenic annotations and (E) universal chromatin state annotations.

After MLE, probes with mean β_mC_ or β_hmC_ < 0.01 across all samples were removed due to increased variability and decreased interpretability of β values at such low levels, as well as to remove the issue of zero inflation for 5hmC β values. We then produced three data sets: one with all probes where β_mC_ > 0.01 (714,427 probes), one with all probes where β_hmC_ > 0.01 (587,091), and one with only probes common to the two sets, where β_mC_ > 0.01 and β_hmC_ > 0.01 (587,065). Of these, zero probes had β_hmC_ data only, and 28,658 had β_mC_ data only.

*Differential testing for differentially methylated cytosines:* The *gamlss* (Generalized Additive Models for Location, Scale, and Shape) R package (version 5.4-22) was used to test for interaction differentially methylated cytosines (iDMCs) as a site where there is a shift in the balance between 5-mC and 5-hmC, as previously described (Supplementary File 3) ^20, 33^. Briefly, the mixed effects model treats 5-mC and 5-hmC as “repeated” measures of a single outcome variable (DNA modification), and a random effect for ID accounts for the correlation between 5-mC and 5-hmC at a CpG site. Meanwhile, a “DNA modification*Experimental Condition” interaction term is used to determine if 5-mC and 5-hmC differ in their response to the experimental condition. PMI was included as a covariate because it was identified as a significant principal component by SVD analysis in *ChAMP*. Age was not included as samples were age-matched, and age was not identified as a significant variable by SVD analysis. Sex was not included because only male samples were included. An FDR < 0.05 was used as the cutoff for significance, and annotation of significant differential probes was performed using the Illumina EPIC array manifests. QQ plots generated using the R package *QCEWAS* (version 1.2.3) and *ggplot2*, respectively, show that these models are appropriately modeling the data, with lambda = 1.22 for interaction modeling.

*Annotation of interaction DMCs:* Gene IDs corresponding to each iDMC were extracted from the EPIC array manifest provided by Illumina (v1.0 B5). Annotation of universal chromatin states was also performed using the *annotatr* R package (version 1.28.0) and adding custom full stack ChromHMM chromatin states for *hg38* to the annotation cache ^34^. For specific candidate loci, brain-specific imputed ChromHMM annotations for cingulate gyrus (E069) and SN (E074) were used; ChromHMM annotations are not available for parietal cortex ^34^. Since this study utilized neuronally enriched nuclei, neuronal expression of iDMC-containing genes was verified using the Allen Brain Cell Atlas ^35^. Specific loci were also compared to a database of imprint control regions (ICR) ^36^.

*Predicting enhancer targets:* iDMCs annotated to weak enhancers, active enhancers, and transcribed enhancers based on ChromHMM chromatin state annotations were input to GREAT to predict target genes of these iDMC-containing enhancers ^37^. The basal plus extension method for association of genes was used, with curated regulatory domains included.

*Gene ontology pathway enrichment:* Gene ontology (GO) term enrichment testing and pathway analysis were performed using a combined list of unique iDMC-containing genes and targets of iDMC-containing enhancer regions using the ClueGO application in Cytoscape (version 3.10.1) ^38, 39^. “Groups” was selected as the visual style, and the GO Biological Process (GOBP) term and the GO Cellular Component term were selected. Network specificity was set to “Medium”, with the GO Tree Interval minimum set at 3 and maximum at 8. Only terms with at least 3 genes and a Bonferroni-corrected p-value < 0.05 were included in pathway visualizations. The connectivity score (Kappa) was set at 0.4, and default GO Term Grouping settings were used in all analyses.

*Protein-protein interaction networks:* The same list of genes was also used for protein-protein interaction network analysis with STRING (version 12.0) using a minimum required interaction score = 0.7 and all other default parameters ^40^. The minimum required confidence level was high >= 0.7.

## Results

### Identification of iDMCs

We identified 108 iDMCs with significant shifts in the proportions of 5mC and 5hmC associated with PD in DNA isolated from an enriched neuronal population derived from parietal cortex (FDR < 0.05) (Figure 1C; Supplementary File 4). Consistent with the known distribution of 5hmC, the majority of iDMCs were found in gene bodies and are located mainly in transcriptionally active chromatin, consistent with their location within gene bodies, and active enhancers (Figure 1D,E).

As shown in Figure 1C, more iDMCs have a negative than a positive β coefficient (76 vs 32, respectively). In this output, a positive β coefficient indicates a relative decrease in 5mC and an increase in 5hmC, as illustrated by visualizing raw BS β values and MLE-estimated β values for the iDMC with the largest positive β coefficient (Figure 2A). In contrast, a negative interaction term indicates a relative increase in 5mC and a decrease in 5hmC, as illustrated by iDMC with the smallest negative β coefficient (Figure 2B).

**Figure 2:**
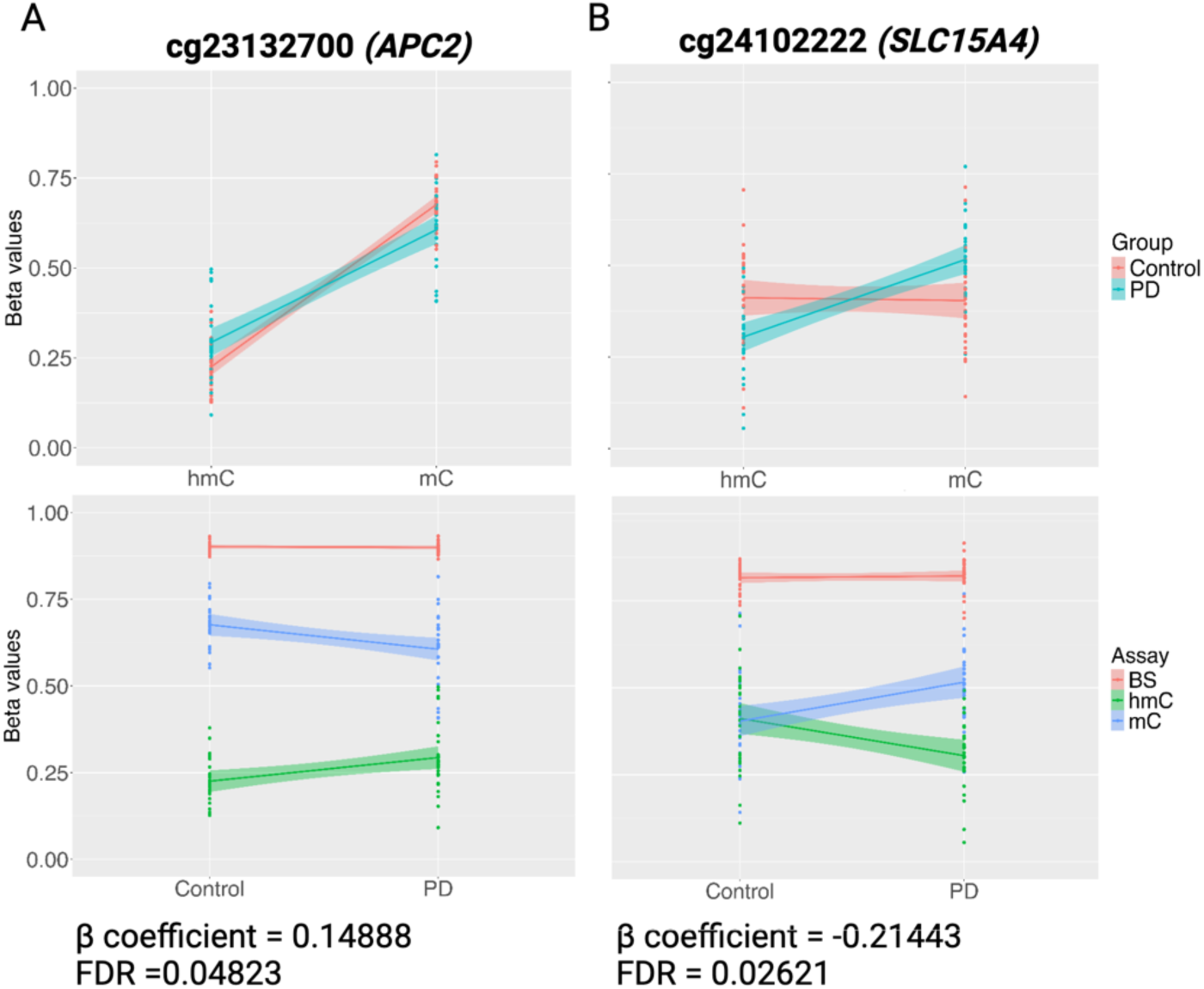
β-values of selected PD-associated DMCs. MLE-corrected βmC and βhmC values graphed by modification (top) and raw BS β values with MLE-corrected βmC and βhmC values graphed by disease status (bottom) for (A) the iDMC with the most positive interaction term and (B) the iDMC with the most negative interaction term (B). The corresponding beta coefficient/interaction term and FDR value are indicated for each iDMC.

### Annotation of iDMCs

Of the 108 iDMCs, 71 were found within genes and were annotated to 83 genes, (Supplementary File 5). Using GREAT, an additional 67 genes were identified as potential targets of iDMC-containing enhancer regions (Supplementary File 6). Of these, 16 were also identified as iDMC-containing genes. Together, 134 unique genes were identified (Supplementary File 7).

### Overlap with previous EWAS studies

Next, we compared these results with our BS-only results and other PD EWAS studies. First, data from our previous publication were reanalyzed. In the previous analysis, intergenic probes were removed prior to differential analysis, but here intergenic probes were included to allow for chromatin annotation of all regions ^19^. While most of the DMCs, DMRs, and ∼95% of genes identified were also identified in the original analysis, some new DMCs and DMRs were significant here, and some were no longer significant (Table 2). The results of this updated analysis were used to compare interaction modeling to BS-only analysis. Annotated DMCs and DMRs from this BS-only analysis are included in Supplementary Files 8 and 9, respectively, and code is provided in Supplementary File 10. Of the 83 iDMC genes identified here, only 7 iDMC-containing genes were also identified by our BS-only analysis (*AGAP1*, *CACNA1H*, *COX19*, *LMF1*, *PMFBP1*, *RAB32* and *TFAP2A*), albeit at different cytosines. Thus, these two methods identified largely unique sets of genes.

**Table 2:** Summary of comparisons between the current study and previous PD EWAS in brain tissue. *Sex not specified, Abbreviations: SN, substantia nigra; PFC, prefrontal cortex; CTX, cortex; FC, frontal cortex; DMV, dorsal motor nucleus of the vagus; CG, cingulate gyrus; PC, parietal cortex

|  | 5hmC | 5mC and 5hmC (BS) |  |  |  |  |  | Paired BS/oxBS |  |
| --- | --- | --- | --- | --- | --- | --- | --- | --- | --- |
|  | Marshall 2020 | Kia 2021 |  | Masliah 2013 | Young 2019 |  |  | Kochmanski 2022 | Current study |
| Control (M/F) | 23* | not specified |  | 6 (2/4) | 41 (all male) |  |  | 49 (29/20) | 29 (all male) |
| PD (M/F) | 20* | 134* |  | 5 (5/0) | 38 (all male) |  |  | 50 (33/17) | 27 (all male) |
| Region | PFC | SN | CTX | FC | DMV | CG | SN | PC | PC |
| Method | hMe-DIP | BS-450K |  | BS-450K | BS-450K/EPIC |  |  | BS-EPIC | BS/oxBS-EPIC |
| # genes | 5157 | 154 | 125 | 155 | 203 | 119 | 1466 | 547 | 83 |
| Overlap with BS | 126 | 5 | 9 | 12 | 20 | 14 | 68 | - | 7 |
| Overlap with interaction | 17 | 0 | 2 | 2 | 3 | 1 | 20 | 7 | - |

Next, we compared the list of genes in this study to recent brain-specific EWAS studies for PD, including ours, for which data was provided ^18, 41–43^. When comparing the current study with 4 recent studies and our BS-only analysis, the most frequently identified genes across studies were *AGAP1, C10orf71, CACNA1H,* and *RAB32* (Supplementary File 11). Of the 4 recent studies, one specifically measured 5hmC by hMeDIP-seq; the others used BS conversion and did not differentiate between 5mC and 5hmC. Finally, 7 genes (*AGAP1, APC2, GNAS, ELANE, POLR2E, ZNF341,* and *WWOX*) were also identified in our two-hit mouse model of increased PD susceptibility ^44, 45^.

### Gene ontology enrichment analysis

By gene ontology enrichment analysis for biological process, 36 genes were enriched in 16 GO terms within 8 GO term groups, including terms related to cell-cell adhesion, multiple cellular development pathways, and chemokine signaling (Table 3). Enrichment analysis for cellular compartment identified 3 enriched GO Terms: catenin complex, catalytic step 2 spliceosome, and anchored compartment of plasma membrane (Table 4).

**Table 3:** Enriched GO Terms and GO Term Groups of identified genes based on the GO Biological Process term. Results of ClueGO gene ontology enrichment analysis and significant GO terms are shown (p < 0.05). GO terms are grouped when they share >50% of their genes. “Associated Genes” shows all genes that map to each GO Term Group.

| GO Groups | GO Terms | Associated Genes |
| --- | --- | --- |
| Group0 | chemokine-mediated signaling pathway | CCR5, CCRL2, STK39, TFF2 |
| Group1 | calcium ion import | CACNA1H, PDGFB, TRPV3 |
| Group2 | neuromuscular process controlling balance | CDH23, GAA, JPH3 |
| Group3 | cell-cell adhesion via plasma-membrane adhesion molecules, homophilic cell adhesion via plasma membrane adhesion molecules | CDH23, FAT3, PCDHGA1, PCDHGA2, PCDHGA3, PCDHGA4, PCDHGA5, PCDHGA6, PCDHGA7, PCDHGA8, PCDHGB1, PCDHGB2, PCDHGB3, PCDHGB4 |
| Group4 | endoderm formation, endodermal cell differentiation | COL5A1, HSBP1, LAMA3 |
| Group5 | phosphatidic acid metabolic process, phosphatidic acid biosynthetic process | AGPAT4, LIPC, PLD1 |
| Group6 | autonomic nervous system development, cranial nerve development | GBX2, PHOX2A, TFAP2A, DRGX |
| Group7 | cellular response to nutrient, response to vitamin D, cellular response to vitamin, vitamin D receptor signaling pathway, cellular response to vitamin D | KANK2, PIM1, RXRA |

**Table 4:** Enriched GO Terms and GO Term Groups of identified genes based on the GO Cellular Component term. Results of ClueGO gene ontology enrichment analysis and significant GO terms are shown (p < 0.05). GO terms are grouped when they share >50% of their genes. “Associated Genes” shows all genes that map to each GO Term Group.

| GO Groups | GO Terms | Associated Genes |
| --- | --- | --- |
| Group0 | catenin complex | APC2, CDH23, PCDHGB4 |
| Group1 | anchored component of plasma membrane | EEPD1, RTN4RL1, TEX101 |
| Group2 | catalytic step 2 spliceosome | DDX23, DHX35, EIF4A3, RBM44 |

### Protein-protein interaction networks

Protein-protein interaction (PPI) networks were generated in STRING and are shown in Figure 3. The largest enriched GO term group and corresponding PPI network contain multiple genes in the protocadherin gamma gene cluster involved in cell-cell adhesion via plasma-membrane adhesion molecules. These protocadherin gamma genes are located in a cluster and are all annotated to a single cytosine, resulting in a large number of genes represented (Table 5, Supplementary File 4). This cytosine is located in a 5′ UTR of *PCDHGA8* but an intron in other *PCDHGA* and *PCDHGB* genes. This site is annotated in ChromHMM as heterochromatin and is located at the boundary with a bivalent promoter in CG (E069) but not in SN (E074) ^34^.

**Figure 3:**
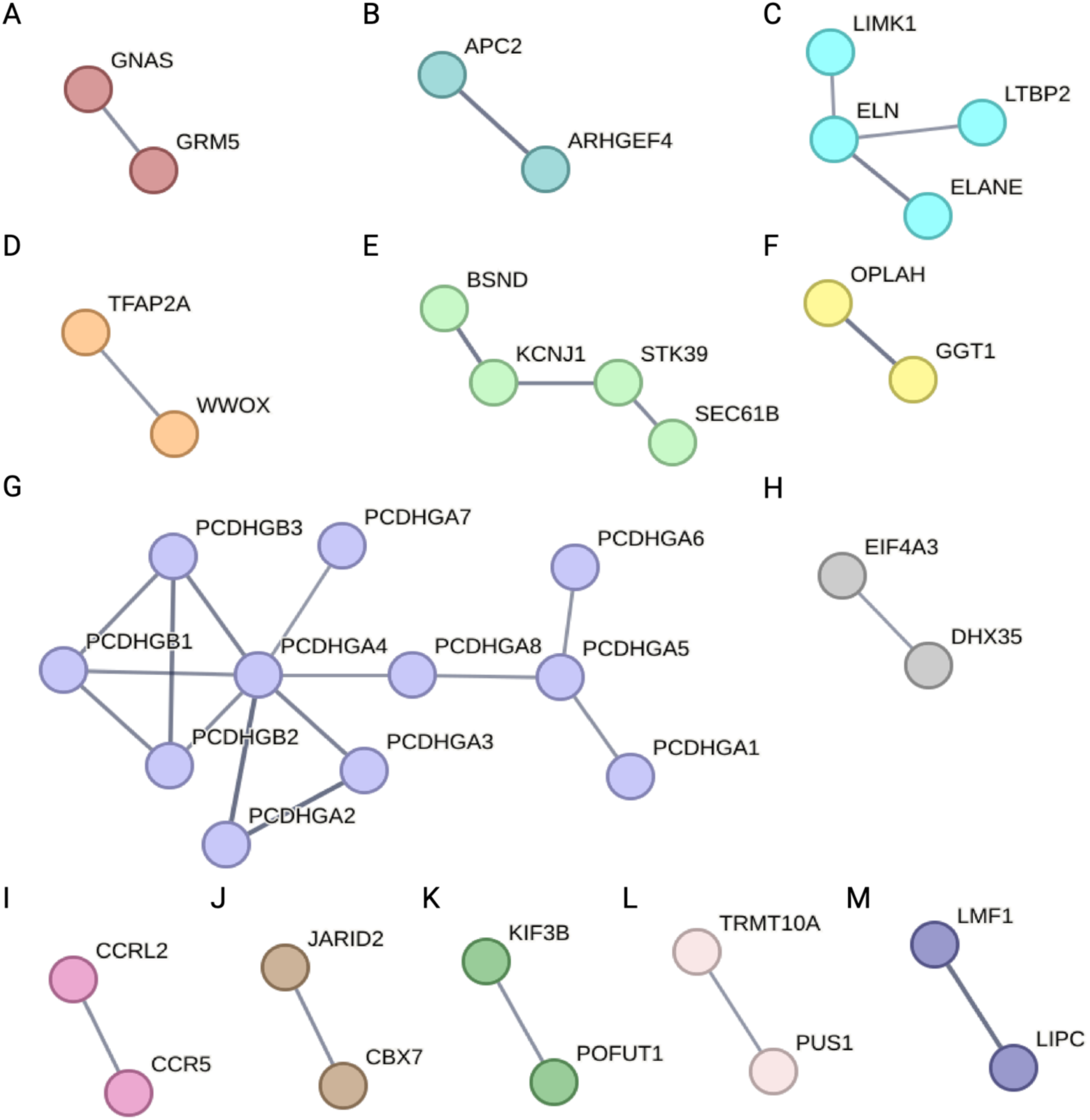
Protein interaction network for iDMC-containing genes and enhancer region target genes. String protein interaction network for 134 unique genes (confidence >= 0.7; >=2 genes per subnetwork. All but 2 genes mapped to the STRING database (MIR7152, LOC102724297). Disconnected nodes in the network are hidden. PPI enrichment value = 2.95 x 10^-7^.

**Table 5:**
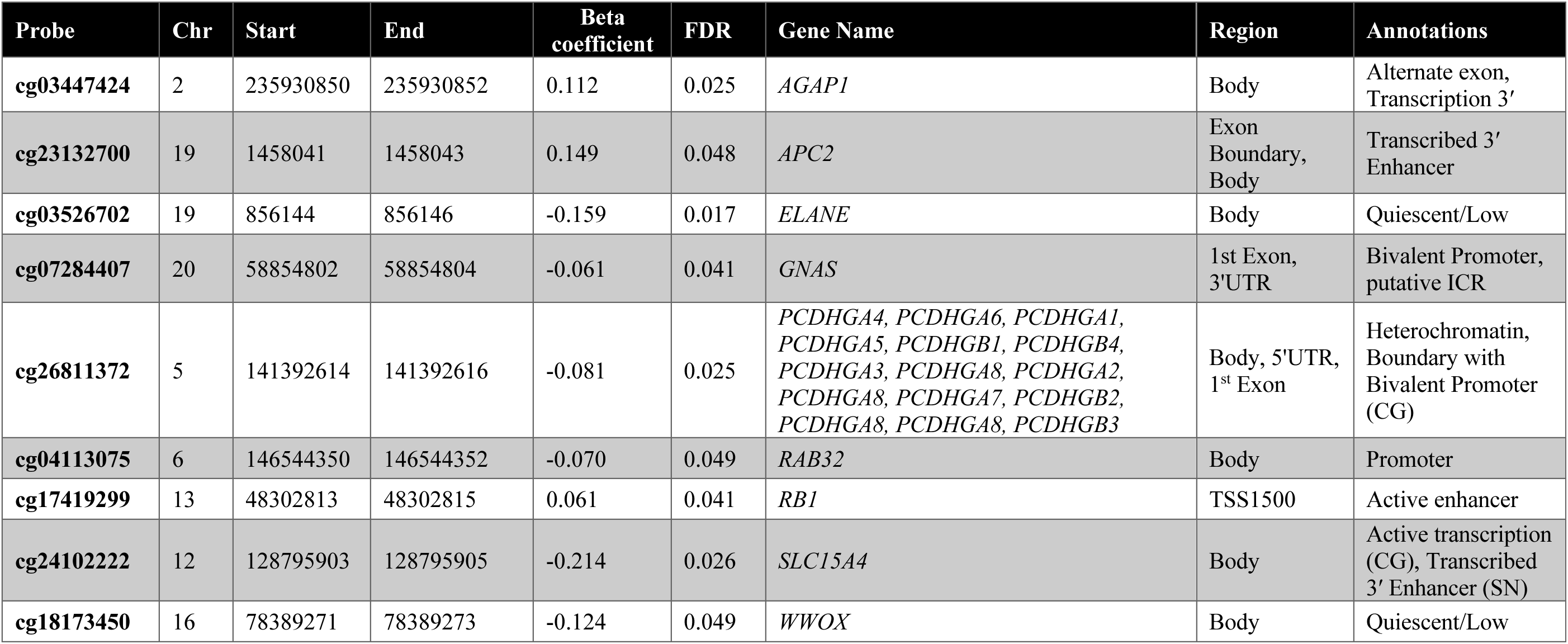
Selected iDMCs of interest. iDMCs highlighted in the text are listed with beta coefficient, FDR, RefSeq Gene Name, region annotation, and brain-specific ChromHMM annotations indicated. Hg38 coordinates are shown. Additional annotations are from brain-specific ChromHMM annotations corresponding to imputed HMM in CG (E069) and SN (E074) and a database of ICRs. When annotations differ between CG and SN, region is indicated.

### Endolysosomal genes

Two of the genes most frequently identified across PD EWAS studies encode the endolysosomal proteins RAB32 and AGAP1 (Supplementary File 11). We identified an iDMC located in a promoter of *RAB32,* showing a relative increase in 5mC and a decrease in 5hmC (Table 5).

Epigenetic regulation of this gene in PD has also been reported in other brain EWAS, in our previous BS-only study, and peripheral immune cells (Table 2, Supplementary File 11) ^18, 19, 43, 46^. Recently, *Rab32* was identified as a causative gene for autosomal dominant PD ^47–51^.

We also identified an iDMC in *AGAP1* within an alternate exon which shows a relative increase in 5hmC and a decrease in 5mC (Table 5). Alternative splicing of this transcript produces at least 3 variants, and this exon is present in at least two of these. Differential modification of this gene was also reported in other brain EWAS and our previous BS-only study (Table 2, Supplementary File 11) ^18, 19, 41^. AGAP1 is an ArfGTase activating protein that has previously been associated with neurodevelopmental disorders, possibly by affecting dendritic spine morphology ^52^. It is a direct regulator of adaptor-related protein complex 3 (AP3) trafficking proteins, and of cytoskeletal remodeling ^53, 54^.

### Genes in neuroinflammatory pathways

The iDMC with the largest negative interaction term annotated to *SLC15A4* and is located within the last intron and annotated as active transcription in CG and transcribed 3ʹ enhancer in SN (Table 5). While SLC15A4 is not shown in the stringent interaction networks generated by STRING in Figure 3, if we allow for interacting proteins and lower the stringency to medium confidence, there are known and potential connections between SLC15A4, NLRP inflammasome proteins, and additional proteins involved in inflammasome activation and neuroinflammation pathways in this dataset: TNFSF11 (TNF superfamily member 11), NFKBID (NF-κB inhibitor delta), IL16 (interleukin 16), CCR5 (C-C chemokine receptor type 5), and CCRL2 (C-C chemokine receptor-like 2). *CCR5* and *CCRL2* are annotated to the GO term “chemokine-mediated signaling pathway” along with *STK39;* these genes appear in two PPI networks (Figure 3E,I, Table 3).

### Imprinted genes

The iDMC-containing imprinted gene *GNAS* is a highly complex imprinted locus with multiple transcripts derived from alternate promoters and 5′ exons, as well as an antisense transcript expressed from the opposite strand that encodes multiple forms of the alpha subunit of the stimulatory G protein (G_⍺_s) ^55^. G_⍺_s acts to couple G protein-coupled receptors for multiple neurotransmitters with their second messenger systems, and proper imprinting plays a critical role in development. The identified iDMC shows a relative increase in 5mC and decrease in 5hmC and is located within a bivalent promoter and putative ICR in an alternate 5′ exon (Table 5, Supplementary File 4) ^36^. In the PPI network, this gene interacts with another iDMC-containing gene *GRM5* (metabotropic glutamate receptor 5, mGluR5), which has been well-studied in the context of PD and L-DOPA-induced dyskinesias, with inconsistent results in clinical trials targeting mGluR5 (Figure 3A) ^56^.

An iDMC annotated to the imprinted gene *RB1* (RB transcriptional corepressor 1) is located within an active enhancer just upstream of the transcription start site for *Rb1* and within an intron of the long non-coding RNA, *RB1* divergent transcript, and shows a relative increase in 5hmC and decrease in 5mC. While Rb1 is extremely well studied in the context of cancer, its role in the nervous system and PD has also been explored, where it is essential for the survival of post-mitotic neurons ^57–59^.

### Genes identified in our model of increased PD susceptibility

Multiple genes identified in this study were also identified in our two-hit mouse model of increased PD susceptibility and form PPI networks with other genes in the current dataset (*AGAP1, APC2, GNAS, ELANE,* and *WWOX*) (Figure 3A-D) ^44, 45^. As discussed above, *GNAS* is an imprinted gene, and it has also been found to be differentially modified in other exposure models ^60^. *APC2* contains the iDMC with the largest positive interaction term (Figure 2A). It encodes the APC2 (APC Regulator Of WNT Signaling Pathway 2), which forms a complex with ARGHEF4 (Rho Guanine Nucleotide Exchange Factor 4) involved in E-cadherin-mediated cell-cell adhesion and regulation of microtubule dynamics with potential functions in axon guidance and dendritic formation during neurodevelopment (Figure 3B) ^61, 62^. APC2 is enriched with the protocadherin gamma genes in the GO terms cell-cell adhesion via plasma-membrane adhesion molecules and catenin complex (Table 5, Figure 3G, Table 3, Table 4).

*ELANE* contains an iDMC within an exon annotated as quiescent/low (Table 5). This gene encodes neutrophil elastase, which has putative connections to ELN (elastin), LIMK1 (LIM domain kinase 1), and LTBP2 (Latent Transforming Growth Factor Beta Binding Protein 2) (Figure 3C). Elastin is one of the main structural components of many tissues, including brain blood vessels, and collectively, these proteins function in creating and maintaining the extracellular matrix (ECM). Within the nervous system, the ECM plays important roles in synaptic plasticity, growth of dendritic spines, and stabilization of synaptic connectivity ^63^. More specifically, recent evidence implicates the degradation of elastin in aging, neuroinflammation, and age-related vascular diseases, but the role of elastin in neurodegenerative disease remains poorly studied ^64^.

*WWOX* contains in iDMC with an intron specific to one *WWOX* transcript variant annotated as quiescent/low (Table 5). This gene encodes the WW domain-containing oxidoreductase, which is known to regulate *TFAP2* (Transcription factor AP-2-alpha) (Figure 3D). While initially identified as a tumor suppressor, *WWOX* plays a role in a wide range of pathways and processes, including neurodevelopment and possibly neurodegeneration ^65^. In addition, *TFAP2* is annotated to GO terms related to nervous system development (Table 4).

## Discussion

Here, we performed an integrated genome-wide analysis of 5mC and 5hmC using our novel application of mixed effects modeling in enriched neuronal nuclei from PD post-mortem parietal cortex samples, which has been used in our lab and others ^20–22^. These PD-associated iDMCs were largely unique from DMCs identified in our previous BS-based EWAS: 7 genes were identified in both studies, 76 only in the paired analysis, and 540 genes only in the BS-only analysis (Supplementary Files 6,7,10) ^19^. Collectively, these data suggest that there are significant PD-associated shifts between 5mC and 5hmC at iDMCs that are not captured by analyzing BS-based data alone (Figure 1C) ^19, 20^. These data suggest that shifts in the balance between DNA modifications may play an important but unrecognized role in PD etiology in both known and novel PD-related genes. While there are many genes of interest to explore in this dataset, here, we highlight a selection of genes based on known functions, gene-ontology enrichment, and protein-protein interaction results. Overall, these results indicate that the inclusion of epigenetic data expands known networks of genes and proteins that may be dysregulated in PD and can identify pathways not previously studied in PD.

We identified multiple genes involved in endolysosomal trafficking and LRRK2-mediated pathways, which are important in PD pathogenesis ^66, 67^. LRRK2 is the most commonly mutated gene in familial PD, and common variants are associated with sporadic PD ^68^. While LRRK2 and these genes are not shown in the stringent interaction networks generated by STRING in Figure 3, if we allow for interacting proteins and lower the stringency, there are known and potential connections between LRRK2 and the following proteins in our dataset: AGAP1, RAB32, RAB41, RADIL, and RAPGEF1. Specifically, Rab32 is a small GTPase that interacts with other PD genes (*LRRK2*, *PINK1*, *VPS35*) that are critical mediators of the endolysosomal sorting pathways known to be involved in PD ^66, 69, 70^. *AGAP1* encodes an ArfGAP and was identified as a differentially expressed gene in peripheral blood samples of fast-and slow-progressing PD patients ^71^. Of particular interest for sporadic PD, defects in AGAP1 function have been proposed to render cells vulnerable to second-hit cytotoxicity and may contribute mechanistically to gene-environment interactions, and LRRK2 can be activated by PD-related toxicants ^68, 72^. The identification of epigenetic regulation of LRRK2-interacting genes suggests that epigenetic regulation of PD risk genes and associated pathways may represent a mechanistic link between genetic and environmental risks for PD.

*Slc15a4* (solute carrier family 15 member 4) contains the iDMC with the largest negative interaction term; SLC15A4 is an amino acid transporter within the endolysosomal membrane involved in the positive regulation of pattern recognition pathways (Figure 2B). While SLC15A4 has primarily been studied in peripheral immune cells, it is expressed in many types of neurons, as verified in the Allen Brain Cell Atlas ^35^. In peripheral immune cells, it is required for trafficking and colocalization of nucleic acid–sensing Toll-like receptors to endolysosmes in conjunction with AP3, and it promotes both inflammasome activity and increased autophagy in response to infection ^73, 74^. Within the brain, the inflammasome is typically thought of in the context of glial cells, but it is also important in neurons, including midbrain neurons ^75–78^.

Epigenetic regulation of these pathways is intriguing, especially given recent interest in the inflammasome as a mediator of gene-environment interactions in PD and ongoing studies of inflammasome inhibitors for PD ^79–82^.

Imprinted genes are highly sensitive to environmental perturbation because their epigenetic marks are not cleared during development and are known to be critical for growth, metabolism, and neuronal function ^83^. Many imprinted genes show distinct patterns of imprinting and expression in the brain compared to other tissues. Environmental disruption of imprinting during development leads to long-term and persistent changes in gene expression in pathways important in the pathogenesis of neurological diseases, providing a potential mechanism by which environmental exposures can impact the risk of late-life disease ^84^. The identified iDMC in *GNAS* is located within bivalent promoter and putative ICR (Table 5). As a result, disruption of imprinting of the *GNAS* locus could lead to changes in imprinting and cell-and tissue-specific transcript expression, affecting development and GPCR-mediated neurotransmitter signaling pathways. The iDMC within the *RB1* locus is not located within the ICR, but epigenetic regulation of the iDMC-containing promoter is known to regulate chromosomal looping and expression of Rb, and disruption of this looping can lead to decreased expression and tumorigenesis ^36, 85^. Thus, it is possible that epigenetic dysregulation in the brain of the chromosomal looping that regulates RB1 expression could lead to cell loss via cell cycle reentry and senescence ^58, 86^.

This dataset is enriched for genes involved cell-cell adhesion by both biological processes and cellular component, including the protocadherin gamma gene cluster (Figure 3G, Table 3, Table 4). The identified locus is in a genomic region annotated as a region-specific promoter (Table 5). Because DNA modification of the protocadherin promoters regulates the differential expression of protocadherin, it is possible that dysregulation of region disrupts the formation and/or maintenance of neuronal circuits and proper synaptic connections ^87, 88^. In our mouse model of increased PD susceptibility, we also identified multiple genes involved in cell-cell adhesion pathway ^44, 45^. Relevant to the current study, a recently proposed hypothesis states that epigenetic dysregulation of this locus plays a role in multiple brain disorders ^89, 90^.

While we assessed neuron-specific DNA modifications associated with PD in a region without widespread neuron loss prior to the onset of pathology within the parietal cortex in an attempt to capture early, pre-degenerative changes, post-mortem studies do not allow for longitudinal analysis of the progressive changes that lead to disease. To model this, we developed a two-hit mouse model of environmentally induced increased PD susceptibility in which developmental exposure to the organochlorine pesticide dieldrin leads to a male-specific exacerbation of neurotoxicity induced by synucleinopathy in the α-synuclein preformed fibril (α-syn PFF) model ^45, 91, 92^. Most recently, we identified dieldrin-induced changes in DNA modifications from birth to 9 months of age in pathways related to early neurodevelopment, dopaminergic neuron differentiation, synaptogenesis, synaptic plasticity, and glial-neuron interactions, consistent with the hypothesis that increased susceptibility to late-onset neurological diseases has origins in development ^93^. The genes and pathways shared between our model and human PD have known functions in neurodevelopment and the establishment and maintenance of synapse and neural circuits, providing insight into mechanisms that may set the stage for increased susceptibility to disease (Table 5, Figure 3C).

While these changes were not assessed in the SN, the region most commonly studied in the context of PD, by using the parietal cortex, which does not have widespread degeneration at this stage of disease, we were able to assess neuron-specific DNA modifications associated with PD in a region without widespread neuron loss. Thus, overall, these data suggest that PD-associated alterations in the epigenetic regulation of these genes may alter gene expression, promoter usage, or isoform expression of these genes and may represent early pre-degenerative events that precede the onset of degeneration.

An additional caveat of these findings is that this study does not address the biological significance of these epigenetic shifts. While shifts between 5mC and 5hmC may potentially impact the binding of proteins that regulate gene expression and/or other epigenetic marks, this study does not examine these functional impacts. However, it does provide multiple avenues for further study of the impact of these changes on gene expression, alternate promoter usage, differential isoform expression, and neuronal function and susceptibility.

It is not surprising that the overlap between our data and these other EWAS was minimal (Table 2, Supplementary File 11). There are major differences in existing PD EWAS related to sample size, sample selection, brain region assessed, methods used to measure DNA modifications, statistical modeling, and inconsistent reporting between studies. For example, one study performed hMe-DIP in prefrontal cortex to assess genome-wide 5hmC, while three BS-based studies used either the EPIC array or the previous 450K array in SN, frontal cortex, the dorsal motor nucleus of the vagus (DMV), and cingulate gyrus (CG) ^18, 41–43^. The methodological differences between these studies complicate the comparison of these studies and underscore the need for rigorous and reproducible methodology and analysis, as discussed in recent chapters from our group and others on rigor and reproducibility in EWAS ^94, 95^.

In addition, it is also important to note that this study included a high level of failed probes and failed samples unique to the oxBS reactions. In contrast, in the BS data, no samples were excluded due to high levels of failed probes and far fewer probes failed. As noted in the methods, we started with 1 µg of input DNA for oxBS because when we used the recommended starting amount of 500 ng, all oxBS probes failed. Together, this suggests that oxBS is much harsher on the DNA that BS alone, requiring high input amounts that may limit the utility of this method as the field moves towards cell type-specific methods and lower input amounts.

## List of supplementary files

Supplementary_File_1_EPIC_QC_MLE.Rmd: RMarkdown file containing code for importing data, performing QC and pre-processing steps, and performing MLE beta value estimation

Supplementary_File_2_meta_data.csv: Meta data file containing phenotypic data and sample sheet data for all samples

Supplementary_File_3_EPIC_DiffMeth.Rmd: RMarkdown file containing code for gamlss interaction modeling to test for differentially methylated and hydroxymethylated CpGs

Supplementary_File_4_Interaction_Male_DMCs_FDR_Filtered_Annotated.csv: Csv file containing significant interaction DMCs with Illumina EPIC manifest and ChromHMM annotations

Supplementary_File_5_Interaction_DMC_Genes_List.txt: Text file containing the list of genes that iDMCs were annotated to

Supplementary_File_6_GREAT_Gene_List.txt: Text file containing the list of genes that are potential targets for iDMC-containing enhancer regions

Supplementary_File_7_iDMC_GREAT_Combined_Gene_List.txt: Text file containing the unique list of genes from Supplementary File 7 and Supplementary File 8

Supplementary_File_8_BSonly_Annotated_DMCs.csv: Csv file containing significant DMCs from reanalysis of previous BS-only analysis

Supplementary_File_9_BSonly_Annotated_DMRs.csv: Csv file containing significant DMRs from reanalysis of previous BS-only analysis

Supplementary_File_10_BSonly_DiffMeth.Rmd: RMarkdown file containing modified code for reanalysis of previous BS-only analysis used to generate Supplementary File 10 and Supplementary File 11

Supplementary_File_11_EWAS_Comparison.xlsx: Excel file containing comparisons between the current study and previous PD EWAS in brain tissue

## Ethics approval and consent to participate

De-identified tissue samples from control (n = 50) and Parkinson’s disease (n = 50) human brain samples were obtained from archival human autopsy specimens provided by the Banner Sun Health Research Institute (BSHRI), using BSHRI’s approved institutional review board (IRB) protocols.

## Data availability

This study was preregistered with Open Science Framework: https://osf.io/z4vbw.

Raw and processed data are available in GEO (GSE267937): https://www.ncbi.nlm.nih.gov/geo/query/acc.cgi?acc=GSE267937

All supplementary material, including additional figures, tables of results, and code used for analyses, are available as supplementary files.

## Author contributions

A.I.B and J.K. designed the study. A.I.B., J.K., N.C.K. and M.A. developed methodology. Experiments were carried out by J.K. and N.C.K. Code was developed, and data was analyzed by N.C.K, J.K, J.I.C, and M.V. J.I.C. and M.V. generated figures and tables. J.I.C, M.V. and A.I.B wrote and edited the manuscript. All authors reviewed the manuscript. A.I.B provided supervision and project administration and acquired funding for this project.

## Acknowledgements

The authors thank the Van Andel Genomics Core, especially Marie Adams, for providing consultation, library preparation, and next-generation sequencing facilities and services (Illumina EPIC array). We are grateful to the Banner Sun Health Research Institute Brain and Body Donation Program of Sun City, Arizona for the provision of human brain tissue. The Banner Sun Health Research Institute Brain and Body Donation Program has been supported by the National Institute of Neurological Disorders and Stroke (U24 NS072026 National Brain and Tissue Resource for Parkinson’s Disease and Related Disorders), the National Institute on Aging (P30 AG19610 Arizona Alzheimer’s Disease Core Center), the Arizona Department of Health Services (contract 211002, Arizona Alzheimer’s Research Center), the Arizona Biomedical Research Commission (contracts 4001, 0011, 05-901 and 1001 to the Arizona Parkinson’s Disease Consortium) and the Michael J. Fox Foundation for Parkinson’s Research.

