## Supplementary material for "Parkinson’s disease-associated alterations in DNA methylation and hydroxymethylation in human brain": Supplementary_File_1_EPIC_QC_MLE.Rmd

````
---
title: "240501_P9_oxBS_QC_MLE_MV"
authors: Nathan Kuhn, Joe Kochmanski, Alison Bernstein, Juliana Choza, Mahek Virani
date: "05/01/2024"
output:
  pdf_document:
    toc: true
  html_document:
    code_folding: hide
    fig_height: 2.5
    fig_width: 3.5
    highlight: tango
    toc: true
    toc_float: true
editor_options:
  chunk_output_type: console
---

```{r setup, include=FALSE}
knitr::opts_chunk$set(echo = TRUE)
```

## Project Summary
This script imports paired BS/oxBS data from the Illumina MethylationEPIC array, performs QC and pre-processing steps, and outputs MLE beta value estimates.

1. Import the data in using minfi and set the annotation to the latest version of the EPIC array annotations (ilm10b4.hg19).
2. Identify probe-level detection p-value and filter probes that have a detection p-value > 0.01 in >10% of samples and probes that failed in any sample.
3. Mask cross-reactive and SNP probes based on Zhou et al. 2017 (PMID:27924034) for EPIC. Documentation and download for Zhou et al. can be found here: http://zwdzwd.github.io/InfiniumAnnotation. 
4. Dye bias correction with ssNoob in minfi.
5. Check for batch effects in ChAMP and adjust with ComBAT.
7. Run ChAMP SVD to determine which variables to include in downstream modeling.
8. Estimate 5-mC and 5-hmC beta values with oxBS.MLE in ENmix.
9. Filter probes with mean 5-mC or 5-hmC beta value < 0.01 

## Data setup
This requires idat files and a sample sheet. The sample sheet must include unique sample names which list the assay used as BS or oxBS within the name (as in "00_09_N_BS"), a column for Assay where 1 indicates BS and 2 indicates oxBS, a column with Subject name such that there are 2 samples/assays per subject, all relevant data about the samples, and all data from running the arrays, including basename relative to this Rmd document.

* Parent Directory
  + This Rmd document
  + meta_data.csv (file containing phenotypic data and sample sheet data).
  + idat (folder contains iScan output files)
  + Manifest (folder contains manifest and/or files from Zhou et al. 2017 (PMID:27924034))
  + Output_QC_MLE_Champ (empty folder for output)
  + Illumina manifest file for EPIC in csv format (download available at: https://support.illumina.com/downloads/infinium-methylationepic-v1-0-product-files.html)

## Load packages
```{r load packages, echo=TRUE, warning=FALSE, message=FALSE, cache=FALSE, results='hide' }
library(posibatch)
library(minfi)
library(IlluminaHumanMethylationEPICanno.ilm10b4.hg19)
library(ChAMP)
library(ENmix)
library(tidyr)
library(plyr)
library(dplyr)
library(quantro)
```

## 1. Import idat files and set annotation
Set annotation to the correct version of the EPIC array in minfi. 
```{r Create RGChannelSet, echo=TRUE, warning=FALSE, message=FALSE, cache=TRUE, results='hold', cache=FALSE}
# Import meta data
meta <- read.csv("./meta_dataV2.csv", colClasses=c("Sentrix_ID"="character"), header = TRUE)
#meta <- meta[, -c(1:15)]
#meta <- subset(meta, select = -c(Sample_Plate, Date))
meta_BS <- filter(meta, Assay_1 == "BS")
meta_oxBS <- filter(meta, Assay_1 == "oxBS")

# Create RGChannelSet objects
rgset_BS <- read.metharray.exp(targets = meta_BS, recursive = TRUE, verbose = TRUE, force = TRUE, extended = TRUE)
rgset_oxBS <- read.metharray.exp(targets = meta_oxBS, recursive = TRUE, verbose = TRUE, force = TRUE, extended = TRUE)

# Set annotation to EPIC
rgset_BS@annotation = c(array='IlluminaHumanMethylationEPIC', annotation='ilm10b4.hg19') #For EPIC array data.
rgset_oxBS@annotation = c(array='IlluminaHumanMethylationEPIC', annotation='ilm10b4.hg19')
```

## 2. Check internal control probes with the ENmix function `plotCtrl`.  
Check internal control probes with the ENmix function `plotCtrl`.  
Review these plots to ensure that the arrays ran correctly. 
```{r Internal control probes, echo=TRUE, warning=FALSE, message=FALSE, cache=TRUE, results='hide'}
dir.create("./Output_QC_MLE_Champ/Control_Probes_BS")
setwd("./Output_QC_MLE_Champ/Control_Probes_BS/")
dim(rgset_BS)
plotCtrl(rgset_BS) 
setwd("../..")

dir.create("./Output_QC_MLE_Champ/Control_Probes_oxBS")
setwd("./Output_QC_MLE_Champ/Control_Probes_oxBS/")
dim(rgset_oxBS)
plotCtrl(rgset_oxBS) 
setwd("../..")
```

## 3. Generate MethylSet without normalization using the minfi function `preprocessRaw`.
Generate MethylSet without normalization using the minfi function `preprocessRaw`.
```{r Generate MethylSet, echo=TRUE, warning=FALSE, message=FALSE, cache=TRUE, results='hold'}

# Set sample names
sampleNames(rgset_BS) <- rgset_BS[["Subject.ID"]]

#Pull out phenotype data
pd_BS <- pData(rgset_BS)

# Generate a MethylSet
mset_BS <- preprocessRaw(rgset_BS)

# Extract the beta values
raw_betas_BS <- getBeta(mset_BS, "Illumina")

sampleNames(rgset_oxBS) <- rgset_oxBS[["Subject.ID"]]

#Pull out phenotype data
pd_oxBS <- pData(rgset_oxBS)

# Generate a MethylSet
mset_oxBS <- preprocessRaw(rgset_oxBS)

# Extract the beta values
raw_betas_oxBS <- getBeta(mset_oxBS, "Illumina")

```

## 4. Filter probes and samples
Generate detection p-values and filter probes with detP > 0.01. 
Then remove probes that failed in > 5% of samples and samples with > 10% probes failed. 
```{r Filter data by detection p-value, echo=TRUE, warning=FALSE, message=FALSE, cache=TRUE}

# Extract detection p-values
detP_BS <- detectionP(rgset_BS)

# Detect low quality probes based on detection p-value
raw_betas_BS[detP_BS >= 0.01] <- NA

# Calculate the proportion of probes that failed the detection p-value threshold
numfail_BS <- matrix(colMeans(is.na(raw_betas_BS)))

# Rename rows and columns
rownames(numfail_BS) <- colnames(detP_BS)
colnames(numfail_BS) <- "Failed CpG Fraction"
numfail_BS

#Repeat for oxBS
detP_oxBS <- detectionP(rgset_oxBS)

# Detect low quality probes based on detection p-value
raw_betas_oxBS[detP_oxBS >= 0.01] <- NA

# Calculate the proportion of probes that failed the detection p-value threshold
numfail_oxBS <- matrix(colMeans(is.na(raw_betas_oxBS)))

# Rename rows and columns
rownames(numfail_oxBS) <- colnames(detP_BS)
colnames(numfail_oxBS) <- "Failed CpG Fraction"
numfail_oxBS

#Check how many and which samples failed in BS and oxBS
write.csv(numfail_BS, "./Output_QC_MLE_Champ/failed_CpG_fraction_BS.csv")
write.csv(numfail_oxBS, "./Output_QC_MLE_Champ/failed_CpG_fraction_oxBS.csv")

SampleCutoff <- 0.1

#Build list of failed samples and subjects that need to be removed
RemovePairs <- c()
RemovePairs <- rownames(numfail_BS)[which(numfail_BS>SampleCutoff)]
RemovePairs <- c(RemovePairs, rownames(numfail_oxBS)[which(numfail_oxBS>SampleCutoff)])
Remove <- unique(RemovePairs)

#Removes subjects from BS data
mset_BS.f <-mset_BS [, !colnames(mset_BS) %in% Remove ]
raw_betas_BS.f <- raw_betas_BS[,!colnames(raw_betas_BS) %in% Remove]
detP_BS.f <- detP_BS[, !colnames(detP_BS) %in% Remove]

meta_BS.f <- meta_BS[!meta_BS$Subject.ID %in% Remove,]
rgset_BS.f <- rgset_BS[, !colnames(rgset_BS) %in% Remove]
pd_BS.f <- pd_BS[!pd_BS$Subject.ID %in% Remove,]
rownames(pd_BS.f) <- meta_BS.f$Sample_Name

#Removes subjects from oxBS data
mset_oxBS.f <-mset_oxBS [, !colnames(mset_oxBS) %in% Remove ]
raw_betas_oxBS.f <- raw_betas_oxBS[,!colnames(raw_betas_oxBS) %in% Remove]
detP_oxBS.f <- detP_oxBS[, !colnames(detP_oxBS) %in% Remove]

meta_oxBS.f <- meta_oxBS[!meta_oxBS$Subject.ID %in% Remove,]
rgset_oxBS.f <- rgset_oxBS[, !colnames(rgset_oxBS) %in% Remove]
pd_oxBS.f <- pd_oxBS[!pd_oxBS$Subject.ID %in% Remove,]
rownames(pd_oxBS.f) <- meta_oxBS.f$Sample_Name

ProbeCutoff <- 0.05
# Remove probes that failed in > 5% of samples change raw_beta.r to raw_betas, will filter out bad data later. 

#BS remove probes and prep matrices for merging
mset_BS.r <- mset_BS.f[rowSums(is.na(raw_betas_BS.f)) <= ProbeCutoff * ncol(detP_BS.f), ]

raw_betas_BS.r <- raw_betas_BS.f[rowSums(is.na(raw_betas_BS.f)) <= ProbeCutoff * ncol(detP_BS.f), ]
colnames(raw_betas_BS.r) <- meta_BS.f$Sample_Name
raw_betas_BS.rt <- as.data.frame(raw_betas_BS.r)
raw_betas_BS.rt$names <- rownames(raw_betas_BS.rt)

detP_BS.r <- detP_BS.f[rowSums(is.na(raw_betas_BS.f)) <= ProbeCutoff * ncol(detP_BS.f),]
colnames(detP_BS.r) <- meta_BS.f$Sample_Name
detP_BS.rt <- as.data.frame(detP_BS.r)
detP_BS.rt$names <- rownames(detP_BS.rt)

#oxBS remove failed probes and prep matrices for merging
mset_oxBS.r <- mset_oxBS.f[rowSums(is.na(raw_betas_oxBS.f)) <= ProbeCutoff * ncol(detP_oxBS.f), ]

raw_betas_oxBS.r <- raw_betas_oxBS.f[rowSums(is.na(raw_betas_oxBS.f)) <= ProbeCutoff * ncol(detP_oxBS.f), ]
colnames(raw_betas_oxBS.r) <- meta_oxBS.f$Sample_Name
raw_betas_oxBS.rt <- as.data.frame(raw_betas_oxBS.r)
raw_betas_oxBS.rt$names <- rownames(raw_betas_oxBS.rt)

detP_oxBS.r <- detP_oxBS.f[rowSums(is.na(raw_betas_oxBS.f)) <= ProbeCutoff * ncol(detP_oxBS.f),]
colnames(detP_oxBS.r) <- meta_oxBS.f$Sample_Name
detP_oxBS.rt <- as.data.frame(detP_oxBS.r)
detP_oxBS.rt$names <- rownames(detP_oxBS.rt)

#Merge BS and oxBS data sets and convert back to matrix
mset.f <- combineArrays(mset_oxBS.r, mset_BS.r)
raw_betas.f <- inner_join(raw_betas_oxBS.rt, raw_betas_BS.rt, by="names")
rownames(raw_betas.f) <- raw_betas.f$names
raw_betas.f$names <- NULL
raw_betas.f <- data.matrix(raw_betas.f)
detP.f <- inner_join(detP_oxBS.rt, detP_BS.rt, by="names")
rownames(detP.f) <- detP.f$names
detP.f$names <- NULL
detP.f <- data.matrix(detP.f)

#Combine meta and rgset files
rgset.f <- minfi::combine(rgset_oxBS.f, rgset_BS.f)
pd.f <- rbind(pd_BS.f, pd_oxBS.f)
meta.f <- bind_rows(meta_BS.f, meta_oxBS.f)

```

## 5. Mask probes
Select probe-lists from Zhou for EPIC.
```{r Mask probes male, echo=TRUE, warning=FALSE, message=FALSE, cache=TRUE}
# Zhou (for EPIC) - Documentation is found here: http://zwdzwd.github.io/InfiniumAnnotation

# Load files
EPIC.manifest <- readRDS("./Manifest/EPIC.hg19.manifest.rds")
head(EPIC.manifest)
EPIC_manifest_hg19 <- as.data.frame(EPIC.manifest)

# Select probes to mask (see documentation here: http://zwdzwd.github.io/InfiniumAnnotation)
maskname <- rownames(EPIC_manifest_hg19)[which(EPIC_manifest_hg19$MASK_general == TRUE)] #99360 probes to mask.

# Filter mset and raw_betas to remove probes
mset.fm <- mset.f[!featureNames(mset.f) %in% maskname, ]
raw_betas.fm <- raw_betas.f[!rownames(raw_betas.f) %in% maskname, ]

```

## 6. Adjust beta-values
Fix Beta values that are either less than or equal to 0 or greater than or equal to 1.
```{r Fix Beta values male, echo=TRUE, warning=FALSE, message=FALSE, cache=TRUE}
if (min(raw_betas.fm, na.rm = TRUE) <= 0)
  raw_betas.fm[raw_betas.fm <= 0] <- min(raw_betas.fm[raw_betas.fm > 0])

if (max(raw_betas.fm, na.rm = TRUE) >= 1)
  raw_betas.fm[raw_betas.fm >= 1] <- max(raw_betas.fm[raw_betas.fm < 1])
```

## 7. Extract raw data and run QC in ChAMP
```{r Extract raw data and QC male, echo=TRUE, warning=FALSE, message=FALSE, cache=TRUE}

#Get the intensity values and detection p-values using minfi to feed to QC in ChAMP
intensity <- minfi::getMeth(mset.fm) + minfi::getUnmeth(mset.fm)
detP.fm <- detP.f[which(row.names(detP.f) %in% row.names(raw_betas.fm)), ]

#Compile the data into a list object to feed into ChAMP QC 
preprocessed.raw.data <- list(mset = mset.fm, rgSet = rgset.f, pd = pd.f, intensity = intensity, beta = raw_betas.fm, detP = detP.fm)

#Run QC with ChAMP
dir.create("./Output_QC_MLE_Champ/ChAMP_Raw")
champ.QC(beta = preprocessed.raw.data$beta, pheno = pd.f$Disease, resultsDir = "./Output_QC_MLE_Champ/ChAMP_Raw")

#Save for easy loading if required
save(mset.fm, raw_betas.fm, detP.fm, intensity, rgset.f, pd.f, meta.f, file = "./Output_QC_MLE_Champ/mset_rgset_rawbetas.RData")

```

## 8. Dye bias correction with ssNoob and run QC
Perform dye bias correction for beta values with ssNoob on probes remaining after filtering and masking. We do not need to separate BS and oxBS at this step since we are using single sample Noob.
```{r Dye bias correction male, echo=TRUE, warning=FALSE, message=FALSE, cache=TRUE, results='hide'}

#Load if required 
load("./Output_QC_MLE_Champ/mset_rgset_rawbetas.RData")

gc()

# Perform dye bias correction for beta values with ssNoob on probes remaining after filtering and masking. We can leave BS and oxBS together because this is single sample Noob. 
mset.n <- preprocessNoob(rgset.f, dyeMethod = "single")[rownames(raw_betas.fm), ]

# Set sample names
sampleNames(mset.n) = mset.n[["Sample_Name"]]

# Extract data from mset
betas.n <- getBeta(mset.n, "Illumina")

#Compile the data into a list object to feed into ChAMP QC
dir.create("./Output_QC_MLE_Champ/ChAMP_ssNoob")
ssNoob.data <- list(mset = mset.n, rgSet = rgset.f, pd = pd.f, intensity = intensity, beta = betas.n, detP = detP.fm)

# Run QC in ChAMP after normalization
champ.QC(beta = ssNoob.data$beta, pheno = pd.f$Disease, resultsDir = "./Output_QC_MLE_Champ/ChAMP_ssNoob/")
```

## 9. Separate data by assay
```{r Separate beta and intensity values and phenotypic data by assay, echo=TRUE, warning=FALSE, message=FALSE, cache=TRUE}

bs.m <- as.character(meta.f$Sample_Name[which(meta.f$Assay == 1)])
betas.n.bs <- betas.n[,bs.m]

ox.m <- as.character(meta.f$Sample_Name[which(meta.f$Assay == 2)])
betas.n.ox <- as.data.frame(betas.n[,ox.m])

colnames(intensity) <- meta.f$Sample_Name
intensity.bs <- as.data.frame(intensity[,bs.m])
intensity.ox <- as.data.frame(intensity[,ox.m])

pd.bs <- pd.f[which(pd.f@listData$Assay == "1") ,]
pd.ox <- pd.f[which(pd.f$Assay == "2") ,]

```

## 10. Run ChAMP SVD
Run SVD to determine which variables to include in downstream modeling.
```{r Run ChAMP on raw BS beta values, echo=TRUE, warning=FALSE, message=FALSE, cache=TRUE, fig.width=7, fig.height=5}
champ.SVD(beta = betas.n.bs %>% as.data.frame(), pd = pd.bs, resultsDir = "./Output_QC_MLE_Champ/CHAMP_SVD_BS/")
champ.SVD(beta = betas.n.ox %>% as.data.frame(), pd = pd.ox, resultsDir = "./Output_QC_MLE_Champ/CHAMP_SVD_OX/")
```

## 11. Positional/Batch correction using ComBat in the posibatch R package
```{r Correct for positional/batch effects in beta values using ComBat, echo=TRUE, warning=FALSE, message=FALSE, cache=TRUE}

#Due to batch effects seen in the output of ChAMP SVD for both BS and oxBS, perform positional/batch correction using ComBat in the posibatch R package for both BS and oxBS

#Create sentrix positional ID data frame
meta.f$sampleNames <- paste(meta.f$Sentrix_ID, meta.f$Array, sep = "_") #Add another sentrix ID for sub-directories
meta.m.f <- filter(meta.f, Assay == 1)
meta.m.sentrix <- meta.m.f[, c("Sample_ID", "sampleNames", "Batch")]
names(meta.m.sentrix)[names(meta.m.sentrix) == "Sample_ID"] <- "sampleID"
names(meta.m.sentrix)[names(meta.m.sentrix) == "Batch"] <- "batches"
names(meta.m.sentrix)[names(meta.m.sentrix) == "sampleNames"] <- "sampleNames"
head(meta.m.sentrix)

#Establish data:
dat.m <-as.matrix(betas.n.bs, row.names = 1)
Sentrix.m <- meta.m.sentrix

#Run positional/batch correction
results.m <- posibatches(dat.m, Sentrix.m, batch=TRUE, par.prior=TRUE, prior.plots=FALSE, mean.only.posi=FALSE, mean.only.batch=FALSE)
bs.fp <- as.data.frame(results.m)

#Check Champ SVD results for posibatch-corrected beta values:
champ.SVD(beta = bs.fp, pd = meta.m.f, resultsDir = "./Output_QC_MLE_Champ/CHAMP_SVD_BS_posibatch/")

#oxBS
#Create sentrix positional ID data frame
meta.f$sampleNames <- paste(meta.f$Sentrix_ID, meta.f$Array, sep = "_") #Add another sentrix ID for sub-directories
meta.h.f <- filter(meta.f, Assay == 2)
meta.h.sentrix <- meta.h.f[, c("Sample_ID", "sampleNames", "Batch")]
names(meta.h.sentrix)[names(meta.h.sentrix) == "Sample_ID"] <- "sampleID"
names(meta.h.sentrix)[names(meta.h.sentrix) == "Batch"] <- "batches"
names(meta.h.sentrix)[names(meta.h.sentrix) == "sampleNames"] <- "sampleNames"
head(meta.h.sentrix)

#Establish data:
dat.h <-as.matrix(betas.n.ox, row.names = 1)
Sentrix.h <- meta.h.sentrix

#Run positional/batch correction
results.h <- posibatches(dat.h, Sentrix.h, batch=TRUE, par.prior=TRUE, prior.plots=FALSE, mean.only.posi=FALSE, mean.only.batch=FALSE)
oxbs.fp <- as.data.frame(results.h)

#Check Champ SVD results for posibatch-corrected beta values:
champ.SVD(beta = oxbs.fp, pd = meta.h.f, resultsDir = "./Output_QC_MLE_Champ/CHAMP_SVD_oxBS_posibatch/")

```

## 12. Remove female samples from downstream analysis
```{r}

#metadata
meta_m<-meta.f[which(meta.f$Sex != 2),]

#beta values
bs.fp_m <- bs.fp[, colnames(bs.fp) %in% c(meta_m$Sample_Name)]
oxbs.fp_m <- oxbs.fp[, colnames(oxbs.fp) %in% c(meta_m$Sample_Name)]

#pd
pd_m <- pd.f[rownames(pd.f) %in% c(meta_m$Sample_Name), ]

```

## 13. Remove outlier from previous study 
```{r Outlier, echo=TRUE, warning=FALSE, message=FALSE, cache=TRUE, results='hold'}

previous_study_meta_data_outlier_removed <- read.csv("./meta_data_outlier_removed.csv", colClasses=c("Sentrix_ID"="character"), header = TRUE) #establish list of sample IDs from previous study
previous_study_filtered_sample_IDs <- previous_study_meta_data_outlier_removed$Sample_ID #establish list of sample IDs from previous study

meta_m_final <- meta_m[meta_m$Sample_ID  %in% previous_study_filtered_sample_IDs ,]

# Remove failed samples from beta values 
bs.fp_m_final <- bs.fp_m[, colnames(bs.fp_m) %in% c(meta_m_final$Sample_Name)]
oxbs.fp_m_final <- oxbs.fp_m[, colnames(oxbs.fp_m) %in% c(meta_m_final$Sample_Name)]

# Remove failed samples from intensity 
intensity.bs_m_final <- intensity.bs[, colnames(intensity.bs) %in% c(meta_m_final$Sample_Name)]
intensity.ox_m_final <- intensity.ox[, colnames(intensity.ox) %in% c(meta_m_final$Sample_Name)]

```

## 14. Run ChAMP SVD
Run SVD to determine which variables to include in downstream modeling.
```{r Run ChAMP on raw BS beta values, echo=TRUE, warning=FALSE, message=FALSE, cache=TRUE, fig.width=7, fig.height=5}

pd.bs <- pd_m[which(pd_m@listData$Assay == "1" & pd_m$Sample_Name  %in% meta_m_final$Sample_Name) ,]
pd.ox <- pd_m[which(pd_m$Assay == "2" & pd_m$Sample_Name  %in% meta_m_final$Sample_Name) ,]

champ.SVD(beta = bs.fp_m_final %>% as.data.frame(), pd = pd.bs, resultsDir = "./Output_QC_MLE_Champ/CHAMP_SVD_BS_outlier_rem/")
champ.SVD(beta = oxbs.fp_m_final %>% as.data.frame(), pd = pd.ox, resultsDir = "./Output_QC_MLE_Champ/CHAMP_SVD_OX_outlier_rem/")

save(meta_m_final, bs.fp_m_final, oxbs.fp_m_final, file = "./Output_QC_MLE_Champ/pre-MLE-data.RData") #Save pre-MLE raw BS and oxBS beta values after all QC steps to use for data visualization

```

## 15. oxBS.MLE
oxBS.MLE requires colnames to match for beta and intensity value objects. This section will depend on how your samples and subjects are named, but is simple if you created a subject ID assigned to each pair in your sample sheet/meta data.
```{r Prep for oxBS MLE in ENmix, echo=TRUE, warning=FALSE, message=FALSE, cache=TRUE}

# Set column names to be the same, using mc.fp rather then betas.n.bs since ran batch correction prior 
colnames(bs.fp_m_final) <- pd.bs@listData$Sample_ID[colnames(bs.fp_m_final) == pd.bs@listData$Sample_Name]
colnames(oxbs.fp_m_final) <- pd.ox@listData$Sample_ID[colnames(oxbs.fp_m_final) == pd.ox@listData$Sample_Name]

colnames(intensity.bs_m_final) <- pd.bs@listData$Sample_ID[colnames(intensity.bs_m_final) == pd.bs@listData$Sample_Name]
colnames(intensity.ox_m_final) <- pd.ox@listData$Sample_ID[colnames(intensity.ox_m_final) == pd.ox@listData$Sample_Name]

```

```{r Run oxBS MLE, echo=TRUE, warning=FALSE, message=FALSE, cache=TRUE}
 
MLE <- oxBS.MLE(beta.BS = as.matrix(bs.fp_m_final), beta.oxBS = as.matrix(oxbs.fp_m_final), N.BS = as.matrix(intensity.bs_m_final), N.oxBS = as.matrix(intensity.ox_m_final))

# Combine  matrices into dataframe
mc_df <- as.data.frame(MLE$`5mC`)
hmc_df <- as.data.frame(MLE$`5hmC`)

#Associates mc values with bs and hmc values with oxbs. While they are functionally identical, important to keep this consistent down the line.
bs.m_final <- bs.m[bs.m %in% meta_m_final$Sample_Name]
ox.m_final <- ox.m[ox.m %in% meta_m_final$Sample_Name]

colnames(mc_df) <- bs.m_final
colnames(hmc_df) <- ox.m_final

#Save final version of meta data used in downstream analysis
write.csv(meta_m_final, "./meta_data_postQC_maleOnly_outlier_rem_Champ_cleaned.csv")

```

## 16. Filter data and prep data for modeling
Filter probes with beta value < 0.01 for 5-hmC and/or 5-mC prior to modeling.
Prep data for downstream modeling. This includes and optional step to chunk the data for multiple node processing. The size and number of these chunks will depend on number of probes and nodes available.
```{r Filtering oxBS.MLE output, echo=TRUE, warning=FALSE, message=FALSE, cache=TRUE}

# Create columns of means
mc_df$mean_5mC <- rowMeans(mc_df, na.rm=TRUE)
hmc_df$mean_5hmC <- rowMeans(hmc_df, na.rm=TRUE)

# Establish probe variable using rownames
mc_df$probe <- rownames(mc_df)
hmc_df$probe <- rownames(hmc_df)

# Filter 5-hmC and 5-mC data to remove probes with mean beta value < 0.01
mc.f <- filter(mc_df, mean_5mC > 0.01)
hmc.f <- filter(hmc_df, mean_5hmC > 0.01)


# Remove Na, possible introduced by MLE
mc.f <- mc.f[complete.cases(mc.f),]
hmc.f <- hmc.f[complete.cases(hmc.f),]
 
```

```{r Prep data for modeling, echo=TRUE, warning=FALSE, message=FALSE, cache=TRUE}

# Merge 5-mC and 5-hmC data for remaining probes and transpose dataframe for downstream processing
bvals <- inner_join(mc.f,hmc.f, by = "probe") 
row.names(bvals) <- bvals$probe 

# Remove extra columns
bvals$mean_5mC <- NULL
bvals$mean_5hmC <- NULL
bvals.complete <- bvals[complete.cases(bvals),]
bvals.complete$probe<-NULL
mc.f$mean_5mC<- NULL
mc.f$probe <- NULL
hmc.f$probe <- NULL
hmc.f$mean_5hmC <- NULL

#Save final versions to use for differential methylation AND hydroxymethylation, also save MLE-corrected beta values to use for data visualization
save(bvals.complete, mc.f, hmc.f, meta_m_final, file = "./Output_QC_MLE_Champ/postQC_mc_hmc_meta.RData")

#Make density plot of beta values, uses quantro package
matdensity(object = as.matrix(bvals.complete), groupFactor = meta_m_final$Disease, xlab = "beta values",
ylab = "density")

```
````
