## Supplementary material for "Parkinson’s disease-associated alterations in DNA methylation and hydroxymethylation in human brain": Supplementary_File_3_EPIC_DiffMeth.Rmd

````
---
title: "240501_P9_oxBS_DiffMeth_MV"
authors: "Nathan Kuhn, Joe Kochmanski, Alison Bernstein, Juliana Choza, Mahek Virani"
date: "05/01/2024"
output: html_document
---
## Project Summary
This script tests for differentially methylated AND hydroxymethylated CpGs using paired gamlss model with interaction term.
1. Test for differentially methylated or hydroxymethylated CpGs using paired gamlss model.
2. Annotate probes using Illumina EPIC manifest.

## Data setup
This script requires the output of EPIC_QC_MLE.Rmd.

* Parent Directory
  + This Rmd document
  + Output_QC_MLE_Champ folder containing output of EPIQ_QC_MLE.Rmd (Output_QC_MLE_Champ/postQC_mc_hmc_meta.RData)
  + Illumina manifest file for EPIC in csv format (download available at: https://support.illumina.com/downloads/infinium-methylationepic-v1-0-product-files.html)
  + Output_DiffMeth (empty folder for output)

## Load packages
```{r load packages, echo=TRUE, warning=FALSE, message=FALSE, cache=FALSE, results='hide' }
library(IlluminaHumanMethylationEPICanno.ilm10b4.hg19)
library(tidyr)
library(plyr)
library(dplyr)
library(gamlss)
library(parallel)
library(ggplot2)
library(quantro)
library(reshape2)
library(annotatr)
library(tidyverse)
```

```{r load data from prev Rmarkdown}
load("./Output_QC_MLE_Champ/postQC_mc_hmc_meta.RData")
```

## 1. Beta value adjustment
```{r adjust beta values for modeling male}
# Transpose for regression
bvals.t <- as.data.frame(t(bvals.complete))
  if (min(bvals.t, na.rm = TRUE) <= 0)
    bvals.t[bvals.t <= 0] <- min(bvals.t[bvals.t > 0])
  
  if (max(bvals.t, na.rm = TRUE) >= 1)
    bvals.t[bvals.t >= 1] <- max(bvals.t[bvals.t < 1])

save(bvals.t, file = "./Output_DiffMeth/bvals_adjusted.RData")
```

## 2. Establish modeling variables
```{r Establish meta data modeling variables}

# Set up appropriate covariates
PMI.m <- meta_m_final$PMI

# Create predictor variables from metadata for downstream modeling; these will vary based on the research question.
disease.m <- meta_m_final$Disease
DNA_mod_cat.m <- meta_m_final$Assay 

# Length of all variables should be the same as the number of sample IDs:
# Build paired ID as a random effect 
pairedID <- nrow(meta_m_final)/2
randeff.m <- as.factor(c(1:pairedID,1:pairedID)) 
```

## 3. Interaction modeling
User will need to specify which variable corresponds to the interaction term, as this may change based on how many covariates are included in the model.
```{r Beta regression gamlss}

numCores <- parallel::detectCores(logical=TRUE) #Detect number of cores available for forking processes.

#Establish function for mixed effects beta regression with a logit link function in gamlss R package:

Ilm450k.beta.fit.full <- function(x, PMI, randeff) {
  x <- as.data.frame(x)
  x$randeff <- randeff
  x$PMI <- PMI
  
# Define data; this will vary depending on sample size of study.
data <- x[1:nrow(meta_m_final),]

# Fit model
none.fit <- gamlss(x ~ disease.m*DNA_mod_cat.m + PMI, random = ~1 | randeff, data = data, sigma.formula = ~ disease.m*DNA_mod_cat.m + PMI, family = NO2, trace = F)
  
# Extract the estimate, se, t-value and p-value for beta coefficient of disease status*DNA modification category (interaction term)
none.fit.sum <- as.data.frame(summary(none.fit))[5,c(1:4)] 
  #Note: May need to change the [7,c(1:4)], depending on how many terms are in the model.
  return(none.fit.sum)
}
 fit_m <- vector(mode='list', length=ncol(bvals.t))
 
# Hush wrapper function used to prevent it from spitting out every single result into the console. Does not otherwise affect the code in any way. The results are entered into a file in the working directory named NUL which will be updated in realtime, so you can use it to check if the function is working and how far along it is.
 
hush=function(code){
  sink("NUL") # use /dev/null in UNIX
  tmp = code
  sink()
  return(tmp)
}

# Run the model. Note that mc.cores here is not platform-agnostic; this code is for a Mac. mc.cores can only be set to 1 on Windows. This code uses parallel::detectCores earlier to identify the number of available cores; set it to 1 on Windows.
#chunk <- bvals.t[,c(1:500)] 
#fitchunk <- vector(mode='list', length=ncol(chunk)) #For testing purposes

fit_m <-  hush(mclapply(as.data.frame(bvals.t), function(x) Ilm450k.beta.fit.full(x, PMI.m, randeff.m), mc.preschedule = F, mc.cores = numCores))

fit_m_with_PMI <- fit_m

save(fit_m_with_PMI, file = "./Output_DiffMeth/interaction_gamlss_fit_m_with_PMI.RData")

```

## 4. Multiple testing correction using false discovery rate
User can set appropriate FDR cutoff and standard error cutoffs.
```{r Use p.adjust function to perform FDR analysis on p-values}

# Create data frames of modeling results for easy sorting:
fit_md <- ldply(fit_m_with_PMI, data.frame) 

# Pull out p-values from modeling results data frame:
p1_m <- fit_md$Pr...t..

# Adjust p-values using Benjamini-Hochberg FDR adjustment method in p.adjust function
fdr_m <- p.adjust(p1_m, method = "BH", n = length(p1_m)) 

# Append the FDR values to the dataset.
fit_md$fdr_m <- fdr_m

save(fit_md, file = "./Output_DiffMeth/interaction_gamlss_fit_md_with_PMI_pre_se_pre_fdr.RData") #Save all resulting probes to use for data visualization

# Filter down to probes with standard error less than 10 to remove any probes with high standard errors. These would be poorly modeled in this analysis.
fit_m.se <- fit_md[(fit_md$Std..Error < 10), ] 

fit_m.se_with_PMI <- fit_m.se

write.csv(fit_m.se_with_PMI, "./Output_DiffMeth/Interaction_Male_DMCs_ALL_with_PMI.csv")

# Filter and sort by FDR value (e.g. FDR<0.05):
fit_m.se.fdr <- fit_m.se[(fit_m.se$fdr_m < 0.05), ]
fit_m.se.fdr.sort <- fit_m.se.fdr[order(fit_m.se.fdr$fdr_m),] 

write.csv(fit_m.se.fdr.sort, "./Output_DiffMeth/Interaction_Male_DMCs_FDR_with_PMI.csv")

```

## 5. Annotate probes
```{r Annotate probes}

# Read in manifest (specify correct file for 450K or EPIC), remove headers, and label probe ID as "probe".
Illumina.EPIC.manifest  <- read.csv("./Manifest/infinium-methylationepic-v-1-0-b5-manifest-file.csv", skip = 7)
colnames(Illumina.EPIC.manifest)[colnames(Illumina.EPIC.manifest) == "Name"]<- "probe"

myvars <- c("probe", "CHR", "MAPINFO", "UCSC_RefGene_Name", "UCSC_RefGene_Accession", "UCSC_RefGene_Group","DMR", "CHR_hg38", "Start_hg38", "End_hg38", "Strand_hg38")

#Limit manifest to only variables of interest (see code directly above):
Illumina.EPIC.manifest.lim <- Illumina.EPIC.manifest[myvars]

# Rename probe ID as "probe" in modeling output
colnames(fit_m.se.fdr.sort)[colnames(fit_m.se.fdr.sort) == ".id"] <- "probe"

#Merge data frames of gamlss interaction term model outputs with the EPIC manifest by "probe" variable:
fit_m_p.merged <- merge(fit_m.se.fdr.sort,Illumina.EPIC.manifest.lim,by=c("probe"))

write.csv(fit_m_p.merged, "./Output_DiffMeth/Interaction_Male_DMCs_FDR_with_PMI_Annotated.csv")

```

```{r}

#Also create custom annotation for full stack ChromHMM chromatin states for human (hg38):
#Human full-stack chromatin state annotation in hg38 assembly downloaded from: https://github.com/ernstlab/full_stack_ChromHMM_annotations
chromatin_db_file = "./hg38_genome_100_segments.bed.gz"
extraCols1 = c(symbol='character')
read_annotations(con = chromatin_db_file, genome = 'hg38', name = 'chromatin',  extraCols = extraCols1, format='bed')

#Double-check that custom annotation track has been successfully added to the mm10
print(annotatr_cache$get('hg38_custom_chromatin'))

```

```{r}

DMC.results <- read.csv("./Output_DiffMeth/Interaction_Male_DMCs_FDR_with_PMI_Annotated.csv")

```

```{r}

DMC.results$chr <- DMC.results$CHR_hg38

DMC.results$start <- DMC.results$Start_hg38-1

DMC.results$end <- DMC.results$End_hg38-2 #based on format of hg38 coordinates in the manifest file

DMC.results.start.col.rm <- dplyr::select(DMC.results, c("chr", "start", "end", "Estimate", "Pr...t..", "fdr_m", "probe"))

#Remove top row of variable names, then save file as .txt file. File should be in standard BED3 format plus any additional columns of sample information.
#This file can then be read in by annotatr to annotate gene regions

#Example for annotatr - BED6+3 format file in the following format:
#chr7 130238539 130238540 -11.041447 2.411176e-28 2.500655e-23 hypo
#chr4 142234769 142234770 -10.707064 9.430525e-27 4.890246e-22 hypo

#Columns 4, 5, 6, and 7 refer to test statistic, p-value, FDR,
#and direction of change, respectively.

#Note that the start column is set to -1 compared to the end column; this 
#ensures that the final form of the data will have identical start and end values
#aftering reading in the data using annotatr.

write.table(DMC.results.start.col.rm, "./Output_DiffMeth/Interaction_Male_DMCs_FDR_with_PMI_BED_format.txt", quote = FALSE, row.names = FALSE, col.names = FALSE, sep = "\t")

#Set working directory and read in bed file
dmc_file = "./Output_DiffMeth/Interaction_Male_DMCs_FDR_with_PMI_BED_format.txt"

#Define and rename extra columns in BED file:
extraCols = c(Estimate = 'numeric', Pr...t.. = 'numeric', fdr_m = 'numeric', probe = 'character')
dmc_regions = read_regions(con = dmc_file, extraCols = extraCols, genome = 'hg38',
                             format = 'bed')
head(dmc_regions)

#Build annotations for hg38 based on variable of interest:
readr::local_edition(1) #avoid bad value vroom error: https://github.com/tidyverse/vroom/issues/519#top
builtin_annotations()
annots_exp = c('hg38_custom_chromatin')
annotations_exp = build_annotations(genome = 'hg38', annotations = annots_exp)

#Annotate the regions:
dmc_annots_exp = annotate_regions(
  regions = dmc_regions,
  annotations = annotations_exp,
  ignore.strand = TRUE)

# Note: annotatr output is a special type of dataframe using the dplyr package.
# By default, dplyr::tbl_df objects have nice printing properties, but it
# hides extra columns that would ordinarily wrap. You can see them all with:
head(as.data.frame(dmc_annots_exp))

#Write the gene annotations to a tab-delimited csv file:

write.csv(as.data.frame(dmc_annots_exp),file="./Output_DiffMeth/Interaction_Male_DMCs_FDR_with_PMI_BED_format_chromHMM_annotated.csv",quote=F, row.names = FALSE)

```

```{r}

#merge manifest and chromHMM annotations

chromHMM.annots <- as.data.frame(dmc_annots_exp)

manifest.chromHMM.merge <- merge(DMC.results, chromHMM.annots, by = c("probe", "Estimate", "Pr...t..", "fdr_m"), all = TRUE)

manifest.chromHMM.merge$X <- NULL
manifest.chromHMM.merge$chr <- NULL
manifest.chromHMM.merge$start.x <- NULL
manifest.chromHMM.merge$end.x <- NULL 
manifest.chromHMM.merge <- rename(manifest.chromHMM.merge,c("start"="start.y"))
manifest.chromHMM.merge <- rename(manifest.chromHMM.merge,c("end"="end.y"))

write.csv(manifest.chromHMM.merge, file="./Output_DiffMeth/Interaction_Male_DMCs_FDR_with_PMI_Manifest_ChromHMM_Annotated.csv",quote=F, row.names = FALSE)

```

```{r}

#select iDMCs annotated to enhancer regions for GREAT analysis

enhancer.chromHMM.annotations <- manifest.chromHMM.merge[grep("Enh", manifest.chromHMM.merge$annot.symbol), ]

enhancer.chromHMM.annotations.annot.range <- dplyr::select(enhancer.chromHMM.annotations, seqnames, annot.start, annot.end, probe) 

write.table(enhancer.chromHMM.annotations.annot.range, sep = "\t", quote = FALSE, col.names = FALSE, row.names = FALSE, "./rGREAT_Enhancer_Annotations/enhancer_annotations_annot_range_bed.bed")

```

```{r}

normal_interaction_hist <- ggplot(fit_md, aes(x=p1_m)) + geom_histogram(bins=150, colour = "transparent") + theme(panel.background = element_blank(), panel.grid.major = element_blank(), panel.grid.minor = element_blank(), axis.line = element_blank()) + xlab("p-value") + ylab("count") + scale_x_continuous(breaks = seq(0, 1, 0.05))
ggsave("./Output_DiffMeth/normal_interaction_pval_hist_smallbins.png", normal_interaction_hist, bg="transparent", dpi = 600)

normal_interaction_hist <- ggplot(fit_md, aes(x=fdr_m)) + geom_histogram(bins=150, colour = "transparent") + theme(panel.background = element_blank(), panel.grid.major = element_blank(), panel.grid.minor = element_blank(), axis.line = element_blank()) + xlab("fdr") + ylab("count") + scale_x_continuous(breaks = seq(0, 1, 0.05))
ggsave("./Output_DiffMeth/normal_interaction_pval_hist_smallbins_fdr.png", normal_interaction_hist, bg="transparent", dpi = 600)

normal_interaction_hist <- ggplot(fit_md, aes(x=t.value)) + geom_histogram(bins=150, colour = "transparent") + theme(panel.background = element_blank(), panel.grid.major = element_blank(), panel.grid.minor = element_blank(), axis.line = element_blank()) + xlab("t-value") + ylab("count")
ggsave("./Output_DiffMeth/normal_interaction_pval_hist_t_value.png", normal_interaction_hist, bg="transparent", dpi = 600)

```

```{r}

fit_m.se.fdr.sort.neg <- fit_m.se.fdr.sort[which(fit_m.se.fdr.sort$Estimate < 0),]
fit_m.se.fdr.sort.pos <- fit_m.se.fdr.sort[which(fit_m.se.fdr.sort$Estimate > 0),]

```
````
