## Supplementary material for "Parkinson’s disease-associated alterations in DNA methylation and hydroxymethylation in human brain": Supplementary_File_10_BSonly_DiffMeth.Rmd

````
---
title: "Supplementary_File_8_BSonly_DiffMeth"
author: "Joe Kochmanski, Alison Bernstein, Mahek Virani"
date: "2024-06-11"
output:
  html_document:
    toc: yes
    toc_float: yes
    df_print: paged
    code_folding: hide
    fig_height: 2.5
    fig_width: 3.5
    highlight: tango
  html_notebook:
    code_folding: hide
    fig_height: 2.5
    fig_width: 3.5
    highlight: tango
    toc: yes
    toc_float: yes
editor_options:
  chunk_output_type: console
---
```{r setup, include=FALSE}
library(knitr)
knitr::opts_chunk$set(echo = TRUE, cache.lazy = FALSE)
```

## Project Summary
This script analyzes previously processed beta values from the Illumina EPIC array and returns differential modified cytosines.

## Data setup
This script requires 2 output files from npjPD_Kochmanski_EPICArray.Rmd (meta_data_cets_outlier_removed.csv and EPIC_betas.RData) and the same folder structure indicated in that Rmd with the indicated addition to the Output folder.
* Parent Directory
  + This Rmd document
  + meta_data_cets_outlier_removed.csv (file containing phenotypic data and sample sheet data, glial content and outliers removed)
  + EPIC_betas.RData (processed beta values)
  + Output_DiffMeth (folder for output)
    - CHAMP_SVD_BS_posibatch (empty folder for ChAMP QC output)
    - Modeling (empty folder for modeling output)
  
## 1. Install and load required packages.
```{r Load packages, echo=TRUE, warning=FALSE, message=FALSE, cache=FALSE, results='hide' }
library(tidyverse)
library(plyr)
#Install package from this paper: https://www.ncbi.nlm.nih.gov/pmc/articles/PMC6021926/
#install.packages("remotes")
#remotes::install_github("ChuanJ/posibatch")
library(posibatch)
library(ChAMP)
library(gamlss)
library(DMRcate)
```

## 2. Import data
Import data if not already loaded into your environment
```{r Import beta values and meta data}
load("./EPIC_betas.RData")
meta <-read.csv("./meta_data_cets_outlier_removed.csv", header = TRUE)
```

## 3. Prep and filter data for modeling
```{r Prep and filter data for modeling, echo=TRUE, warning=FALSE, message=FALSE, cache=TRUE}

# Create columns of means
mc$mean_5mC <- rowMeans(mc, na.rm=TRUE)

# Establish probe variable using rownames
mc$probe <- rownames(mc)

# Remove probes with mean beta value < 0.01 (difficult to model and interpret)
mc.f <- mc[mc$mean_5mC > 0.01,]

# Remove extra columns
mc.f$probe <- NULL
mc.f$mean_5mC <- NULL
```
After filtering for mean beta value < 0.01, `r toString(nrow(mc)-nrow(mc.f))` probes were removed. 
`r toString(nrow(mc.f))` probes remain.

## 4. Positional/Batch correction 
Using ComBat in the posibatch R package to correct for positional and batch effects. Confirm that this correction reduced these effects using ChAMP SVD.
```{r Correct for positional/batch effects in beta values using ComBat, echo=TRUE, warning=FALSE, message=FALSE, cache=TRUE}

#Create sentrix positional ID data frame
meta$sampleNames <- paste(meta$Sentrix_ID, meta$Array, sep = "_") 
meta.sentrix <- meta[, c("Sample_ID", "sampleNames", "Batch")]
names(meta.sentrix)[names(meta.sentrix) == "Sample_ID"] <- "sampleID"
names(meta.sentrix)[names(meta.sentrix) == "Batch"] <- "batches"
names(meta.sentrix)[names(meta.sentrix) == "sampleNames"] <- "sampleNames"

#Establish data:
dat <-as.matrix(mc.f, row.names = 1)
Sentrix <- meta.sentrix

#Run positional/batch correction
results <- posibatches(dat, Sentrix, batch=TRUE, par.prior=TRUE, prior.plots=FALSE, mean.only.posi=FALSE, mean.only.batch=FALSE)
mc.fp <- as.data.frame(results)

#Save the data for easy loading 
save(mc.fp,file = "./Output_DiffMeth/EPIC_betas_positionalcorrection.RData")

#Load previous data
load("./Output_DiffMeth/EPIC_betas_positionalcorrection.RData")

#Run ChAMP SVD results for posibatch-corrected beta values:
champ.SVD(beta = mc.fp, pd = meta, resultsDir = "./Output_DiffMeth/CHAMP_SVD_BS_posibatch/")
```

## 6. Stratify data by sex 
```{r Prep data for modeling, echo=TRUE, warning=FALSE, message=FALSE, cache=TRUE}

#Split meta data by sex
meta.m <- meta[which(meta$Gender != 2),] #62 samples
meta.f <- meta[which(meta$Gender != 1),] #37 samples

#Split beta values by sex
bvals <- mc.fp
bvals.f <- bvals[ , names(bvals) %in% c(meta.f$Sample_ID)]
bvals.m <- bvals[ , names(bvals) %in% c(meta.m$Sample_ID)]
```
There are `r toString(nrow(meta.m))` male samples and `r toString(nrow(meta.f))` female samples.

## 7. Modeling - female data
The modeling steps that follow are for females only. Modeling for males is below.

### Include all probles
```{r include all probes for modeling females, echo=TRUE, warning=FALSE, message=FALSE, cache=TRUE}

#Transpose for regression
bvals.t.f <- as.data.frame(t(bvals.f))

#Save beta values and meta data for female data for future loading:
save(bvals.t.f, meta.f,file = "./Output_DiffMeth/EPIC_betas_Female.RData")

#Load beta values and meta data for female data for future loading:
#load("./Output_DiffMeth/EPIC_betas_female.RData")

#Create small data chunk to test code
#bvals.t.f <- bvals.t.f[,c(1:20000)]

numCores <- parallel::detectCores(logical=TRUE) #Detect number of cores available for forking processes.
library(parallel)

#Chunk the data for multiple node processing if necessary
bvals.t.f.chnk1 <- bvals.t.f[,c(1:100000)]
bvals.t.f.chnk2 <- bvals.t.f[,c(100001:200000)]
bvals.t.f.chnk3 <- bvals.t.f[,c(200001:300000)]
bvals.t.f.chnk4 <- bvals.t.f[,c(300001:400000)]
bvals.t.f.chnk5 <- bvals.t.f[,c(400001:500000)]
bvals.t.f.chnk6 <- bvals.t.f[,c(500001:600000)]
bvals.t.f.chnk7 <- bvals.t.f[,c(600001:700000)]
bvals.t.f.chnk8 <- bvals.t.f[,c(700001:761119)]

```

### Create covariates for GAMLSS modeling
Create covariates from the metadata for downstream modeling. These must be set by the user based on the research question. Here we include glial cell proportion and age. 
```{r Establish meta data modeling variables, echo=TRUE, warning=FALSE, message=FALSE, cache=TRUE}

# Set up appropriate covariates
glial <- meta.f$glial
age <- meta.f$Age

# Create predictor variables from metadata for downstream modeling
disease <- meta.f$Disease

#Note: Length of all variables must be the same as the number of sample IDs.
```

###GAMLSS regression analysis with a Gaussian distribution AND sigma term to account for variance
User will need to specify variables while setting up this function, as this may change based on how many covariates are included in the model.

```{r gamlss}
#Establish function for beta regression models with logit link functions in gamlss R package:
IlmEPIC.sigma.fit.sep <- function(x, glial, age) {
  x <- as.data.frame(x)
  x$glial <- glial
  x$age <- age

  # Define data
  data.5mC <- x[1:(length(disease)),]
  
  # Fit the model for the 5-mC data
  mc.fit <- gamlss(x ~ disease + glial + age, data = data.5mC, sigma.formula = ~ disease+ glial + age, 
                   family = NO2, trace = F)
  
  # Extract the estimate, se, t-value and p-value for beta coefficient (interaction term)
  mc.fit.sum <- as.data.frame(summary(mc.fit))[2,c(1:4)] 
  # Note: May need to change the [2,c(1:4)], depending on order of terms in the model
  
  return(list(model.5mC = mc.fit.sum))
}

# Run model on all probes
#s.fit <- mclapply(as.data.frame(bvals.t.f), function(x) IlmEPIC.sigma.fit.sep(x, glial, age), mc.preschedule = F, mc.cores = 4) 
#Create data frame
#s.fit <- ldply (s.fit, data.frame)
#Save data frame of results
#save(s.fit, file = "./Output_DiffMeth/Modeling/EPIC_Female_fit.RData") 

#Load if modeling results have already been saved
#load("./Output_DiffMeth/Modeling/EPIC_Female_fit.RData"")

# Optional: Run the model on each chunk
s.fit1 <- mclapply(as.data.frame(bvals.t.f.chnk1), function(x) IlmEPIC.sigma.fit.sep(x, glial, age), mc.preschedule = F, mc.cores = numCores) #Model
s.fit.1 <- ldply (s.fit1, data.frame) #Create data frame
save(s.fit.1, file = "./Output_DiffMeth/Modeling/EPIC_Female_fit1.RData") #Save data frame of results

s.fit2 <- mclapply(as.data.frame(bvals.t.f.chnk2), function(x) IlmEPIC.sigma.fit.sep(x, glial, age), mc.preschedule = F, mc.cores = numCores)
s.fit.2 <- ldply (s.fit2, data.frame) #Create data frame
save(s.fit.2, file = "./Output_DiffMeth/Modeling/EPIC_Female_fit2.RData") #Save data frame of results

s.fit3 <- mclapply(as.data.frame(bvals.t.f.chnk3), function(x) IlmEPIC.sigma.fit.sep(x, glial, age), mc.preschedule = F, mc.cores = numCores)
s.fit.3 <- ldply (s.fit3, data.frame) #Create data frame
save(s.fit.3, file = "./Output_DiffMeth/Modeling/EPIC_Female_fit3.RData") #Save data frame of results

s.fit4 <- mclapply(as.data.frame(bvals.t.f.chnk4), function(x) IlmEPIC.sigma.fit.sep(x, glial, age), mc.preschedule = F, mc.cores = numCores)
s.fit.4 <- ldply (s.fit4, data.frame) #Create data frame
save(s.fit.4, file = "./Output_DiffMeth/Modeling/EPIC_Female_fit4.RData") #Save data frame of results

s.fit5 <- mclapply(as.data.frame(bvals.t.f.chnk5), function(x) IlmEPIC.sigma.fit.sep(x, glial, age), mc.preschedule = F, mc.cores = numCores)
s.fit.5 <- ldply (s.fit5, data.frame) #Create data frame
save(s.fit.5, file = "./Output_DiffMeth/Modeling/EPIC_Female_fit5.RData") #Save data frame of results

s.fit6 <- mclapply(as.data.frame(bvals.t.f.chnk6), function(x) IlmEPIC.sigma.fit.sep(x, glial, age), mc.preschedule = F, mc.cores = numCores)
s.fit.6 <- ldply (s.fit6, data.frame) #Create data frame
save(s.fit.6, file = "./Output_DiffMeth/Modeling/EPIC_Female_fit6.RData") #Save data frame of results

s.fit7 <- mclapply(as.data.frame(bvals.t.f.chnk7), function(x) IlmEPIC.sigma.fit.sep(x, glial, age), mc.preschedule = F, mc.cores = numCores)
s.fit.7 <- ldply (s.fit7, data.frame) #Create data frame
save(s.fit.7, file = "./Output_DiffMeth/Modeling/EPIC_Female_fit7.RData") #Save data frame of results

s.fit8 <- mclapply(as.data.frame(bvals.t.f.chnk8), function(x) IlmEPIC.sigma.fit.sep(x, glial, age), mc.preschedule = F, mc.cores = numCores)
s.fit.8 <- ldply (s.fit8, data.frame) #Create data frame
save(s.fit.8, file = "./Output_DiffMeth/Modeling/EPIC_Female_fit8.RData") #Save data frame of results

```

### Combine differential modeling results if data was run in chunks
```{r Consolidate data frames of modeling results, echo=TRUE, warning=FALSE, message=FALSE, cache=TRUE}

#Load saved data chunks
#load("./Output_DiffMeth/Modeling/EPIC_Female_fit1.RData"")
#load("./Output_DiffMeth/Modeling/EPIC_Female_fit2.RData")
#load("./Output_DiffMeth/Modeling/EPIC_Female_fit3.RData"")
#load("./Output_DiffMeth/Modeling/EPIC_Female_fit4.RData")
#load("./Output_DiffMeth/Modeling/EPIC_Female_fit5.RData"")
#load("./Output_DiffMeth/Modeling/EPIC_Female_fit6.RData")

# Bind the data frames together with rbind:
s.fit.df1 <- rbind(s.fit.1,s.fit.2)
s.fit.df2 <- rbind(s.fit.df1,s.fit.3)
s.fit.df3 <- rbind(s.fit.df2,s.fit.4)
s.fit.df4 <- rbind(s.fit.df3,s.fit.5)
s.fit.df5 <- rbind(s.fit.df4,s.fit.6)
s.fit.df6 <- rbind(s.fit.df5, s.fit.7)
s.fit <- rbind(s.fit.df6,s.fit.8)

#Save data frame of all results
save(s.fit, file = "./Output_DiffMeth/Modeling/EPIC_Female_fit.RData") 

#Check dataframe of results
summary(s.fit)

#Check p-value histogram of results. If model was appropriately constructed, there should be a clear peak near zero that then tapers down to a uniform distribution.
hist(s.fit$model.5mC.Pr...t..) 
```
Model has been run for `r toString(nrow(s.fit))` probes. 

Sigma term regression results saved to ./Output_DiffMeth/Modeling/ 

Example output of modeling:  
`r toString(s.fit[1])`

### Apply p-value cutoff to modeling results
User can set appropriate standard error cutoff and FDR, q-value, or p-value cutoffs.
```{r Use p.adjust function to perform FDR analysis on p-values for modeling, echo=TRUE, warning=FALSE, message=FALSE, cache=TRUE}

# Extract p-values from modeling results
p.mc <- s.fit$model.5mC.Pr...t..

#If desired, can adjust p-values using Benjamini-Hochberg FDR adjustment method in p.adjust function
fdr.mc <- p.adjust(p.mc, method = "BH", n = length(p.mc)) 

#Can also generate q-values (analogous to FDR) using the qvalue package:
#BiocManager::install("qvalue")
library(qvalue)
qvalue.mc <- qvalue(p.mc)

#Append the FDR values to the dataset.
s.fit$fdr.5mC <- fdr.mc
s.fit$qvalue <- qvalue.mc$qvalues

#Keep probes with standard error less than 10 to remove probes with high standard errors. These would be poorly modeled in this analysis.
s.fit.se <- s.fit[(s.fit$model.5mC.Std..Error < 10), ] 

#Filter and sort by p-value cutoff (e.g. p<9E-8):
s.fit.se <- s.fit.se[(s.fit.se$model.5mC.Pr...t.. < 0.00000009), ]
s.fit.se.sort <- s.fit.se[order(s.fit.se$model.5mC.Pr...t..),] 

```
After filtering for probes by standard error, `r toString(nrow(s.fit.se))` of `r toString(nrow(s.fit))` probes remain.

After filtering for probes by p-value cutoff (p<9E-8), there are `r toString(nrow(s.fit.se.sort))` significant differentially methylated probes in female samples.

### Annotate differentially methylated probe IDs
User can select annotation details to keep
```{r Annotate probes, echo=TRUE, warning=FALSE, message=FALSE, cache=TRUE}

#Load Illumina EPIC manifest
Illumina.EPIC.manifest <- read.csv("./Manifest/MethylationEPIC_v-1-0_B4.csv", skip=7)

#Create subset of variables of interest to keep in downstream analyses:
myvars <- c("Name", "CHR", "MAPINFO", "UCSC_RefGene_Name", "UCSC_RefGene_Accession", "UCSC_RefGene_Group", "UCSC_CpG_Islands_Name", "Relation_to_UCSC_CpG_Island", "Phantom4_Enhancers", "Phantom5_Enhancers", "DMR","X450k_Enhancer", "HMM_Island", "Regulatory_Feature_Name", "Regulatory_Feature_Group", "DNase_Hypersensitivity_NAME", "DNase_Hypersensitivity_Evidence_Count", "OpenChromatin_NAME", "OpenChromatin_Evidence_Count", "TFBS_NAME", "TFBS_Evidence_Count")

#Limit manifest to only variables of interest (see code directly above):
Illumina.EPIC.manifest.lim <- Illumina.EPIC.manifest[myvars]

# Rename probe ID as "Name" in modeling output
colnames(s.fit.se.sort)[colnames(s.fit.se.sort) == ".id"] <- "Name"

# Merge data frames of gamlss interaction term model outputs with the EPIC manifest by "probe" variable:
s.fit.mc.merged <- merge(s.fit.se.sort,Illumina.EPIC.manifest.lim,by=c("Name"))

#Now sort by P-value again!
s.fit.mc.merged.sort <- s.fit.mc.merged[order(s.fit.mc.merged$model.5mC.Pr...t..),] 

#Save annotated results
write.csv(s.fit.mc.merged.sort, file = "./Output_DiffMeth/EPIC_Female_fit_manifest_annotated.csv")
```
Significant probes have been annotated.

These are the top ten hits.
`r toString head(s.fit.mc.merged.sort,n=10)`

### Run DMRcate to test for DMRs
```{r DMRcate to test for DMRs, echo=TRUE, warning=FALSE, message=FALSE, cache=TRUE}

#Establish design matrix for modeling:
design <- model.matrix(~factor(meta.f$Disease) + meta.f$glial + meta.f$Age)

#Convert to data matrix:
bvals.f.matrix <- data.matrix(bvals.f)

#Run DMRcate annotate function:
myannotation <- cpg.annotate("array", bvals.f.matrix, arraytype = "EPIC", analysis.type="differential", design=design, coef=2)
#By default, the contrast may return no individually significant probes. If so, try increasing the fdr. Alternatively, set #pcutoff manually in dmrcate() to return DMRs, but be warned there is an increased risk of Type I errors.

#Run dmrcate on annotated features:
dmrcoutput <- dmrcate(myannotation, lambda=1000, C=2, pcutoff = 0.05)

#Convert DMR list to a GRanges object, which uses the genome argument to annotate overlapping gene loci.
results.ranges <- extractRanges(dmrcoutput, genome = "hg19")

dmr.ranges.f <- as.data.frame(results.ranges)

#Sort by min smoothed FDR:
dmr.ranges.f.sort <- dmr.ranges.f[order(dmr.ranges.f$min_smoothed_fdr),] 

#Save DMRs as output:
write.csv(dmr.ranges.f.sort, "./Output_DiffMeth/EPIC_Female_DMRs.csv")
```
Significant DMRs have been detected and annotated.  

There are `r toString(nrow(dmr.ranges.f))` DMRs.  

The top ten DMRs are:  
`r toString(head(dmr.ranges.f.sort), n=10)`

## 8. Modeling - male data
The modeling steps that follow are for males only. Modeling for females is above.

### Include all probes
```{r include all probes for modeling males, echo=TRUE, warning=FALSE, message=FALSE, cache=TRUE}

#Transpose for regression
bvals.t.m <- as.data.frame(t(bvals.m))

#Save beta values and meta data for male data for future loading:
save(bvals.t.m, meta.m,file = "./Output_DiffMeth/EPIC_betas_male.RData")

#Load beta values and meta data for male data for future loading:
load("./Output_DiffMeth/EPIC_betas_male.RData")

#Create small data chunk to test code
#bvals.t.m <- bvals.t.m[,c(1:20000)]
```

### Create covariates for GAMLSS modeling
Create covariates from the metadata for downstream modeling. These must be set by the user based on the research question. Here we include glial cell proportion and age. 
```{r Establish meta data modeling variables, echo=TRUE, warning=FALSE, message=FALSE, cache=TRUE}

# Set up appropriate covariates
glial <- meta.m$glial
age <- meta.m$Age

# Create predictor variables from metadata for downstream modeling
disease <- meta.m$Disease

#Note: Length of all variables must be the same as the number of sample IDs.
```

###GAMLSS regression analysis with a Gaussian distribution AND sigma term to account for variance
User will need to specify variables while setting up this function, as this may change based on how many covariates are included in the model.

```{r gamlss}

#Establish function for beta regression models with logit link functions in gamlss R package:
IlmEPIC.sigma.fit.sep <- function(x, glial, age) {
  x <- as.data.frame(x)
  x$glial <- glial
  x$age <- age

  # Define data
  data.5mC <- x[1:(length(disease)),]
  
  # Fit the model for the 5-mC data
  mc.fit <- gamlss(x ~ disease + glial + age, data = data.5mC, sigma.formula = ~ disease+ glial + age, 
                   family = NO2, trace = F)
  
  # Extract the estimate, se, t-value and p-value for beta coefficient (interaction term)
  mc.fit.sum <- as.data.frame(summary(mc.fit))[2,c(1:4)] 
  # Note: May need to change the [2,c(1:4)], depending on order of terms in the model
  
  return(list(model.5mC = mc.fit.sum))
}

numCores <- parallel::detectCores(logical=TRUE) #Detect number of cores available for forking processes.
library(parallel)

#Chunk the data for multiple node processing if necessary
bvals.t.m.chnk1 <- bvals.t.m[,c(1:100000)]
bvals.t.m.chnk2 <- bvals.t.m[,c(100001:200000)]
bvals.t.m.chnk3 <- bvals.t.m[,c(200001:300000)]
bvals.t.m.chnk4 <- bvals.t.m[,c(300001:400000)]
bvals.t.m.chnk5 <- bvals.t.m[,c(400001:500000)]
bvals.t.m.chnk6 <- bvals.t.m[,c(500001:600000)]
bvals.t.m.chnk7 <- bvals.t.m[,c(600001:700000)]
bvals.t.m.chnk8 <- bvals.t.m[,c(700001:761119)]

# Run model on all probes
#s.fit.m <- mclapply(as.data.frame(bvals.t.m), function(x) IlmEPIC.sigma.fit.sep(x, glial, age), mc.preschedule = F, mc.cores = numCores) 
#Create data frame
#s.fit.m <- ldply (s.fit.m, data.frame)
#Save data frame of results
#save(s.fit.m, file = "./Output_DiffMeth/Modeling/EPIC_male_fit.RData") 
```

```{r gamlss}

#Load if modeling results have already been saved
#load("./Output_DiffMeth/Modeling/EPIC_male_fit.RData")

# Optional: Run the model on each chunk
s.fit.m1 <- mclapply(as.data.frame(bvals.t.m.chnk1), function(x) IlmEPIC.sigma.fit.sep(x, glial, age), mc.preschedule = F, mc.cores = numCores) #Model
s.fit.m.1 <- ldply (s.fit.m1, data.frame) #Create data frame
save(s.fit.m.1, file = "./Output_DiffMeth/Modeling/EPIC_male_fit1.RData") #Save data frame of results

s.fit.m2 <- mclapply(as.data.frame(bvals.t.m.chnk2), function(x) IlmEPIC.sigma.fit.sep(x, glial, age), mc.preschedule = F, mc.cores = numCores)
s.fit.m.2 <- ldply (s.fit.m2, data.frame) #Create data frame
save(s.fit.m.2, file = "./Output_DiffMeth/Modeling/EPIC_male_fit2.RData") #Save data frame of results

s.fit.m3 <- mclapply(as.data.frame(bvals.t.m.chnk3), function(x) IlmEPIC.sigma.fit.sep(x, glial, age), mc.preschedule = F, mc.cores = numCores)
s.fit.m.3 <- ldply (s.fit.m3, data.frame) #Create data frame
save(s.fit.m.3, file = "./Output_DiffMeth/Modeling/EPIC_male_fit3.RData") #Save data frame of results

s.fit.m4 <- mclapply(as.data.frame(bvals.t.m.chnk4), function(x) IlmEPIC.sigma.fit.sep(x, glial, age), mc.preschedule = F, mc.cores = numCores)
s.fit.m.4 <- ldply (s.fit.m4, data.frame) #Create data frame
save(s.fit.m.4, file = "./Output_DiffMeth/Modeling/EPIC_male_fit4.RData") #Save data frame of results

s.fit.m5 <- mclapply(as.data.frame(bvals.t.m.chnk5), function(x) IlmEPIC.sigma.fit.sep(x, glial, age), mc.preschedule = F, mc.cores = numCores)
s.fit.m.5 <- ldply (s.fit.m5, data.frame) #Create data frame
save(s.fit.m.5, file = "./Output_DiffMeth/Modeling/EPIC_male_fit5.RData") #Save data frame of results

s.fit.m6 <- mclapply(as.data.frame(bvals.t.m.chnk6), function(x) IlmEPIC.sigma.fit.sep(x, glial, age), mc.preschedule = F, mc.cores = numCores)
s.fit.m.6 <- ldply (s.fit.m6, data.frame) #Create data frame
save(s.fit.m.6, file = "./Output_DiffMeth/Modeling/EPIC_male_fit6.RData") #Save data frame of results

s.fit.m7 <- mclapply(as.data.frame(bvals.t.m.chnk7), function(x) IlmEPIC.sigma.fit.sep(x, glial, age), mc.preschedule = F, mc.cores = numCores)
s.fit.m.7 <- ldply (s.fit.m7, data.frame) #Create data frame
save(s.fit.m.7, file = "./Output_DiffMeth/Modeling/EPIC_male_fit7.RData") #Save data frame of results

s.fit.m8 <- mclapply(as.data.frame(bvals.t.m.chnk8), function(x) IlmEPIC.sigma.fit.sep(x, glial, age), mc.preschedule = F, mc.cores = numCores)
s.fit.m.8 <- ldply (s.fit.m8, data.frame) #Create data frame
save(s.fit.m.8, file = "./Output_DiffMeth/Modeling/EPIC_male_fit8.RData") #Save data frame of results
```

### Combine differential modeling results if data was run in chunks
```{r Consolidate data frames of modeling results, echo=TRUE, warning=FALSE, message=FALSE, cache=TRUE}

#Load saved data chunks
#load("./Output_DiffMeth/Modeling/EPIC_male_fit1.RData")
#load("./Output_DiffMeth/Modeling/EPIC_male_fit2.RData")
#load("./Output_DiffMeth/Modeling/EPIC_male_fit3.RData")
#load("./Output_DiffMeth/Modeling/EPIC_male_fit4.RData")
#load("./Output_DiffMeth/Modeling/EPIC_male_fit5.RData")
#load("./Output_DiffMeth/Modeling/EPIC_male_fit6.RData")
#load("./Output_DiffMeth/Modeling/EPIC_male_fit7.RData")
#load("./Output_DiffMeth/Modeling/EPIC_male_fit8.RData")

# Bind the data frames together with rbind:
s.fit.m.df1 <- rbind(s.fit.m.1,s.fit.m.2)
s.fit.m.df2 <- rbind(s.fit.m.df1,s.fit.m.3)
s.fit.m.df3 <- rbind(s.fit.m.df2,s.fit.m.4)
s.fit.m.df4 <- rbind(s.fit.m.df3,s.fit.m.5)
s.fit.m.df5 <- rbind(s.fit.m.df4,s.fit.m.6)
s.fit.m.df6 <- rbind(s.fit.m.df5,s.fit.m.7)
s.fit.m <- rbind(s.fit.m.df6,s.fit.m.8)

#Save data frame of all results
save(s.fit.m, file = "./Output_DiffMeth/Modeling/EPIC_male_fit.RData") 

#Check dataframe of results
summary(s.fit.m)

#Check p-value histogram of results. If model was appropriately constructed, there should be a clear peak near zero that then tapers down to a uniform distribution.
hist(s.fit.m$model.5mC.Pr...t..) 
```
Model has been run for `r toString(nrow(s.fit.m))` probes. 

Sigma term regression results saved to ./Output_DiffMeth/Modeling/ 

Example output of modeling:  
`r toString(s.fit.m[1])`

### Apply p-value cutoff to modeling results
User can set appropriate standard error cutoff and FDR, q-value, or p-value cutoffs.
```{r Use p.adjust function to perform FDR analysis on p-values for modeling, echo=TRUE, warning=FALSE, message=FALSE, cache=TRUE}

# Extract p-values from modeling results
p.mc.m <- s.fit.m$model.5mC.Pr...t..

#If desired, can adjust p-values using Benjamini-Hochberg FDR adjustment method in p.adjust function
fdr.mc.m <- p.adjust(p.mc.m, method = "BH", n = length(p.mc.m)) 

#Can also generate q-values (analogous to FDR) using the qvalue package:
#BiocManager::install("qvalue")
library(qvalue)
qvalue.mc.m <- qvalue(p.mc.m)

#Append the FDR values to the dataset.
s.fit.m$fdr.5mC <- fdr.mc.m
s.fit.m$qvalue <- qvalue.mc.m$qvalues

#Keep probes with standard error less than 10 to remove probes with high standard errors. These would be poorly modeled in this analysis.
s.fit.m.se <- s.fit.m[(s.fit.m$model.5mC.Std..Error < 10), ] 

#Filter and sort by p-value cutoff (e.g. p<9E-8):
s.fit.m.se <- s.fit.m.se[(s.fit.m.se$model.5mC.Pr...t.. < 0.00000009), ]
s.fit.m.se.sort <- s.fit.m.se[order(s.fit.m.se$model.5mC.Pr...t..),] 

```
After filtering for probes by standard error, `r toString(nrow(s.fit.m.se))` of `r toString(nrow(s.fit.m))` probes remain.

After filtering for probes by p-value cutoff (p<9E-8), there are `r toString(nrow(s.fit.m.se.sort))` significant differentially methylated probes in male samples.

### Annotate differentially methylated probe IDs
User can select annotation details to keep
```{r Annotate probes, echo=TRUE, warning=FALSE, message=FALSE, cache=TRUE}

#Load Illumina EPIC manifest
Illumina.EPIC.manifest <- read.csv("./Manifest/MethylationEPIC_v-1-0_B4.csv", skip=7)

#Create subset of variables of interest to keep in downstream analyses:
myvars <- c("Name", "CHR", "MAPINFO", "UCSC_RefGene_Name", "UCSC_RefGene_Accession", "UCSC_RefGene_Group", "UCSC_CpG_Islands_Name", "Relation_to_UCSC_CpG_Island", "Phantom4_Enhancers", "Phantom5_Enhancers", "DMR","X450k_Enhancer", "HMM_Island", "Regulatory_Feature_Name", "Regulatory_Feature_Group", "DNase_Hypersensitivity_NAME", "DNase_Hypersensitivity_Evidence_Count", "OpenChromatin_NAME", "OpenChromatin_Evidence_Count", "TFBS_NAME", "TFBS_Evidence_Count")

#Limit manifest to only variables of interest (see code directly above):
Illumina.EPIC.manifest.lim <- Illumina.EPIC.manifest[myvars]

# Rename probe ID as "Name" in modeling output
colnames(s.fit.m.se.sort)[colnames(s.fit.m.se.sort) == ".id"] <- "Name"

# Merge data frames of gamlss interaction term model outputs with the EPIC manifest by "probe" variable:
s.fit.m.mc.merged <- merge(s.fit.m.se.sort,Illumina.EPIC.manifest.lim,by=c("Name"))

#Now sort by P-value again!
s.fit.m.mc.merged.sort <- s.fit.m.mc.merged[order(s.fit.m.mc.merged$model.5mC.Pr...t..),] 

#Save annotated results
write.csv(s.fit.m.mc.merged.sort, file = "./Output_DiffMeth/EPIC_male_fit_manifest_annotated.csv") 

```
Significant probes have been annotated.

These are the top ten hits.
`r toString head(s.fit.m.mc.merged.sort,n=10)`

### Run DMRcate to test for DMRs
```{r DMRcate to test for DMRs, echo=TRUE, warning=FALSE, message=FALSE, cache=TRUE}

#Establish design matrix for modeling:
design <- model.matrix(~factor(meta.m$Disease) + meta.m$glial + meta.m$Age)

#Convert to data matrix:
bvals.m.matrix <- data.matrix(bvals.m)

#Run DMRcate annotate function:
myannotation <- cpg.annotate("array", bvals.m.matrix, arraytype = "EPIC", analysis.type="differential", design=design, coef=2)
#By default, the contrast may return no individually significant probes. If so, try increasing the fdr. Alternatively, set #pcutoff manually in dmrcate() to return DMRs, but be warned there is an increased risk of Type I errors.

#Run dmrcate on annotated features:
dmrcoutput <- dmrcate(myannotation, lambda=1000, C=2, pcutoff = 0.05)

#Convert DMR list to a GRanges object, which uses the genome argument to annotate overlapping gene loci.
results.ranges <- extractRanges(dmrcoutput, genome = "hg19")

dmr.ranges.m <- as.data.frame(results.ranges)

#Sort by min smoothed FDR:
dmr.ranges.m.sort <- dmr.ranges.m[order(dmr.ranges.m$min_smoothed_fdr),] 

#Save DMRs as output:
write.csv(dmr.ranges.m.sort, "./Output_DiffMeth/EPIC_male_DMRs.csv")
```
Significant DMRs have been detected and annotated.  

There are `r toString(nrow(dmr.ranges.m))` DMRs.  

The top ten DMRs are:  
`r toString(head(dmr.ranges.m.sort), n=10)`
````
